# Skin mesenchymal niches maintain and protect AML-initiating stem cells

**DOI:** 10.1101/2022.05.20.491183

**Authors:** Lakshmi Sandhow, Huan Cai, Elory Leonard, Pingnan Xiao, Luana Tomaipitinca, Alma Månsson, Makoto Kondo, Xiaoyan Sun, Anne-Sofie Johansson, Karl Tryggvason, Maria Kasper, Marcus Järås, Hong Qian

## Abstract

Leukemia cutis or leukemic cell infiltration in skin is one of the common extramedullary manifestations of acute myeloid leukemia (AML) and signifies a poorer prognosis. However, its pathogenesis and maintenance remain understudied. Here, we report massive AML cell infiltration in the skin in a transplantation-induced MLL-AF9 AML mouse model. These AML cells could regenerate AML post-transplantation. Prospective niche characterization revealed that skin harbored mesenchymal progenitor cells (MPCs) with a similar phenotype as BM mesenchymal stem cells. These skin MPCs protected AML-initiating stem cells (LSCs) from chemotherapy *in vitro* partially via mitochondrial transfer. Furthermore, *Lama4* deletion in skin MPCs promoted AML LSC proliferation and chemoresistance. Importantly, more chemoresistant AML LSCs appeared to be retained in *Lama4*^-/-^ mouse skin post-cytarabine treatment. Our study reveals the characteristics and previously unrecognized roles of skin mesenchymal niches in maintaining and protecting AML LSCs during chemotherapy, meriting future exploration of their impact on AML relapse.

**A 40-word summary**

Sandhow et al have in transplantation-induced AML mouse models demonstrated the leukemia-regenerating capacity of AML cells infiltrated in the skin and the role of skin mesenchymal niches in maintaining/protecting AML cells, providing new insight into the pathology of leukemia cutis.

## Introduction

Leukemia cutis (leukemic cell infiltration in skin) is one of the commonly observed extramedullary manifestations in acute myeloid leukemia (AML) (Bakst et al., 2011; Gouache et al., 2018) It is associated with poor survival (Gouache et al., 2018; Krooks and Weatherall, 2018; Wang et al., 2019). However, our understanding of its pathogenesis is very limited. This has been mainly due to lack of knowledge about the extramedullary niches where AML cells infiltrate and are maintained.

For decades, great efforts have been put to elucidate the contribution of the hematopoietic niche in bone marrow (BM) to normal and malignant hematopoiesis. The BM niche consists of various types of cells including mesenchymal stem cells (MSCs), mesenchymal progenitor cells (MPCs), osteoblasts and endothelial cells. Accumulating evidence has shown that BM MSCs and MPCs play important roles in maintaining normal hematopoiesis and in the development of myeloid malignancies including AML (Hoggatt et al., 2016; Xiao et al., 2018b). Molecular alterations in these niches could lead to malignant transformation of hematopoietic cells (Dong et al., 2016; Raaijmakers et al., 2010; Xiao et al., 2018a). During leukemia development, the leukemic cells can alter the BM niche, which renders the niche to become favorable for leukemic cell growth but harmful for normal hematopoiesis (Hanoun et al., 2014; Jeong et al., 2018; Kim et al., 2015; Schepers et al., 2013; Xiao et al., 2018b). Using a *Cre-LoxP* mouse model allowing for selective deletion of the MSC population, we have shown that loss of MSCs accelerates AML progression (Hanoun et al., 2014; Xiao et al., 2018b). However, these markers are not unique for the mesenchymal cells in BM, thus, it is not clear whether the mesenchymal cell niches in extramedullary organs like skin are also affected, thereby contributing to the AML progression.

Besides BM, extramedullary organs such as adipose tissue have been reported to accommodate hematopoietic stem and progenitor cells (HSPCs) in the stromal vascular fraction (Han et al., 2010). In blast crisis of chronic myeloid leukemia, the gonadal white adipose tissue is infiltrated by leukemic cells that adapt to the extramedullary niche and transform into resistant leukemia-initiating stem cells (LSCs) marked by CD36 expression (Ye et al., 2016).These findings indicate that extramedullary organs may serve as niches for HSPCs and LSCs.

Existing evidence suggests that skin might contain an MSC-like cell population (Kimlin and Virador, 2013; Toma et al., 2001; Vaculik et al., 2012; Viswanathan et al., 2019). However, their biological properties and contribution to hematopoiesis and leukemia remain unexplored. Knowledge of this would be essential for understanding the pathology of extramedullary leukemia in the skin, and would thus provide new insights into identifying new potential therapeutic targets that may be translated into new treatment strategies for AML patients with leukemia cutis.

We here in a transplantation-induced AML mouse model detected massive infiltration of AML cells in the skin. Importantly, these skin AML cells at steady-state or post-chemotherapy could regenerate AML post-secondary transplantation even with a low dose of 10 cells/mouse. Characterization of the AML niche revealed the molecular features and developmental hierarchy of the skin MPCs. The MPCs expressing Early B-cell Factor 2 (Ebf2) contribute to mesenchymal cell turnover in the skin and were reduced post-AML onset. The skin MPCs could maintain and protect AML LSCs, and loss of *Lama4* in skin MPCs promoted AML cell maintenance and resistance to chemotherapy. Altogether, our study revealed the characteristics and a previously unrecognized role of skin MPCs in maintaining AML LSCs during AML development and chemotherapy.

## Results

### The AML cells infiltrated in skin can regenerate AML post-transplantation

We have here studied extramedullary leukemia in skin by using an MLL-AF9^+^ AML cell transplantation-induced AML mouse model (**Figure 1A**). Post-symptomatic AML onset (day 21-30 post-transplantation), we detected the AML cell infiltration in the dorsal skin by fluorescence-activated cell sorting (FACS) (**Figure 1B-1D**) and hematoxylin and eosin staining (**Figure S1A**). The AML frequency was higher than that in the blood, but lower than that in BM and spleen (**Figure 1B-1C**). There was no clear correlation between the AML burden in skin and that in blood (**Figure S1B**). However, interestingly, the AML engraftment level in the skin correlated with a faster AML onset (**Figure 1D**) but, not with the AML burden in blood, pointing to a possible predicting value of AML cells in skin for AML progression. Confocal imaging showed that these AML cells mainly resided at perivascular sites in the skin (**Figure 1E**). The skin total cellularity was not affected (**Figure S1C**).

**Figure 1.**
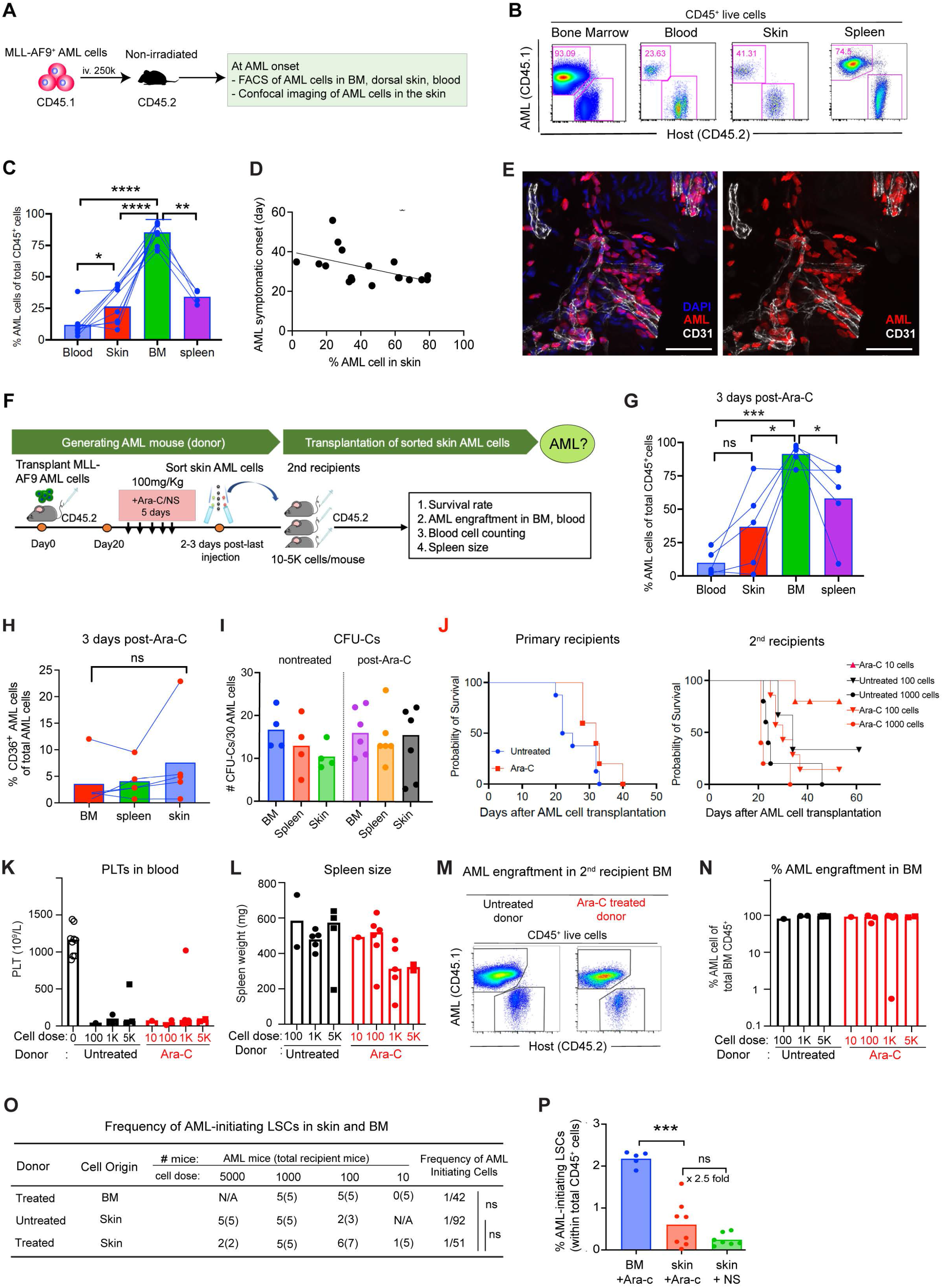
AML cells infiltrated in skin are capable to regenerate AML post- transplantation. (**A**) Strategy to generate MLL-AF9 AML mouse model. MLL-AF9 transduced AML cells expressing CD45.1 were transplanted into non-irradiated CD45.2 C57BL/6 mice. (**B**) FACS profile showing AML engraftment analysis in BM, peripheral blood (PB), spleen and dorsal skin at the end stage of AML. The numbers in the panel are the mean frequencies. (**C**) Proportion of the AML cells within total hematopoietic (CD45^+^) cell in blood, skin, BM and spleen. Data were from 2 independent experiments and each dot represents data from one mouse. The horizontal bars represent mean values. n=4-9 per group, * P<0.05, ** P<0.01, **** P<0.0001, determined by paired *t*-test except the comparison between skin and blood where unequal number of the mice were included. (**D**) The inverse correlation between AML engraftment in the skin with symptomatic AML onset. * P<0.05, determined by Spearman correlation. Each dot represents data from a single recipient mouse, n=18 mice. (**E**) Confocal image showing distribution of MLL-AF9^+^ AML cells expressing DsRed at perivascular sites in dorsal skin. The AML cells were injected after sublethal irradiation of the mice. The endothelial cells were marked by CD31 expression (white). Scale bars are 50 µm. (**F**) Experimental strategy to assess AML-initiating capacity of the AML cells infiltrated in mouse skin by serial transplantation. The primary recipient mice (CD45.2) that developed AML post-AML cell (CD45.1) transplantation were treated with Ara-C (100mg/Kg) or saline (NS, untreated) for 5 days. The residual CD45.1^+^ AML cells from skin were sorted at 2-3 days after the last injection of Ara-C and transplanted into secondary non-irradiated recipient CD45.2 mice at doses of 10, 100, 1000 (1K) and 5000 (5K) cells/mouse. The AML development was monitored by FACS and hematology analyzer Sysmex. The mice were sacrificed when found moribund. (**G**) AML engraftment in blood, skin, BM and spleen of the primary recipients under steady state and 3 days post-Ara-C treatment. Each dot represents data from a single mouse. n=5 per group, * P<0.05, **** P<0.0001, determined by paired *t*-test (**H**) The fraction of CD36^+^ AML cells in the skin, BM and spleen. Each dot represents data from a single mouse. ns, no significant difference, determined by paired *t*-test. (**I**) The frequency of CFU-Cs derived from the residual AML cells in skin, spleen and BM. (**J**) The Kaplan-Meier survival curve of the primary and secondary recipient mice. The curve was generated by Log-rank (Mantel-Cox test). Each dot represents an endpoint when the mice were found dead or sacrificed because of being moribund. Data were from 2 independent experiments. n=5-8 primary recipients treated with Ara-C (n=5) or normal saline (untreated), n=5-7 secondary recipients per group. (**K**) Platelet counts in PB of the secondary recipient mice at the endpoint. Each dot represents data from a single mouse. (**L**) Spleen size of the secondary recipient mice at the endpoint. Each dot represents data from a single mouse. (**M**) FACS profile showing the engraftment of skin-derived AML CD45.1^+^ cells in BM of the secondary recipient mice. (**N**) AML engraftment of the skin-derived AML CD45.1^+^ cells in BM of the secondary recipient mice. Each dot represents data from a single mouse. (**O**) The frequencies of the secondary recipient mice that developed AML after injection of skin-derived AML cells. The frequency of AML-initiating LSCs was determined by Extreme Limiting Dilution analysis (Hu and Smyth, 2009) based on the number of mice that developed AML post-secondary transplantation. ns, no significant difference, determined by *Chi* square test. (**P**) The % of AML-initiating LSCs in BM and skin. The percentages were calculated based on the frequencies of AML-initiating LSCs determined by transplantation and shown in (G) and the % AML cells within total CD45^+^ mononuclear cells in each tissue. Each dot represents data from a single mouse. n=5-8 per group, ns, no significant difference, **** P<0.0001, determined by unpaired *t*-test. See also in Figure S1.

It is not known whether the AML cells infiltrated in skin contain LSCs capable to regenerate leukemia, particularly after chemotherapy. We therefore analyzed the AML cells in the skin 3 days post-5-day treatment with cytarabine (Ara-C)/normal saline (NS) or at the endpoint when the mice became moribund (**Figure 1F**). The AML engraftment in skin remained lower than that in BM and spleen post-Ara-C (**Figure 1G, Figure S1D**), however, there was no clear difference in the fraction of chemoresistant CD36^+^ AML cells and colony-forming unit-in culture (CFU-C) in the AML cells from different tissues (**Figure 1H-1I**).

To functionally assess the frequency of the AML-initiating LSCs in the skin and BM, we sorted the AML cells from the primary recipient mice post-Ara-C treatment (100mg/Kg, daily) and performed secondary transplantation into non-irradiated mice in limiting dilution at doses of 10, 100, 1000 and 5000 cells per mouse (**Figure 1F**). The primary recipients treated with NS showed clear severe symptoms at day 22-33 resulting in ethical endpoint with an estimated median survival of 23.5 days. The median survival of the Ara-C-treated mice was 32 days (**Figure 1J**). The AML cells were sorted at 2-3 days after the last injection of Ara-C or NS. Notably, these AML cells in the skin could generate AML even at a low dose (10-100 cells/mouse) (**Figure 1J-1O**), as shown by the survival (**Figure 1J**), low platelet counts (**Figure 1K**), splenomegaly (**Figure 1L**) and high AML engraftment in the BM at the endpoint (**Figure 1M-1N)**. Extreme Limiting Dilution analysis (Hu and Smyth, 2009) revealed that the frequency of residual AML-initiating LSCs in the skin (1/51) is similar to that in the BM (1/42) post-Ara-C treatment, but seemed to be somewhat higher than that (1/92) from the nontreated skin (**Figure 1O)**. Given the higher AML cell frequency in the BM and spleen, the % of the AML-initiating LSCs and CFU-Cs in BM and spleen became higher than that in the skin (**Figure S1E**, **Figure 1P**). These data demonstrate the leukemia-initiating capacity of the AML cells in the skin tissue under steady-state and after chemotherapy, indicating their potential impact on AML relapse.

### Skin harbors Ebf2^+^ and Ebf2^-^ MPC subsets with similar immunophenotype to BM MSCs

To understand how the leukemic cells were maintained in the skin tissue, we explored the role of the extramedullary microenvironment in the skin using a similar approach as was used for characterizing the BM niche. BM MSCs have been shown to be involved in AML progression in mice (Xiao et al., 2018b). They are defined by a phenotype of lacking expression of CD44, hematopoietic (CD45/TER119) and endothelial (CD31) cell markers, but positive for PDGFRa/CD140a, SCA1 and CD51 (Morikawa et al., 2009; Pinho et al., 2013; Schepers et al., 2013; Xiao et al., 2018a; Xiao et al., 2018b). The primitive MSCs in mouse BM can be identified by Ebf2 expression (Qian et al., 2013) and can generate MSCs that lack Ebf2 (Xiao et al., 2018b). However, the phenotype of native skin MSCs remains unknown. The question is whether the cells with similar features exist in skin and play a role in the AML cell maintenance under steady-state and following chemotherapy. To answer this, we first characterized mesenchymal stem and progenitors in the skin by using the *Ebf2-Egfp* reporter mice. Similar to BM, a fraction of CD45^-^TER119^-^CD31^-^ skin stromal cells expressed Ebf2 (**Figure 2A**). The Ebf2^+^ cells accounted for about 2% and 5% of total live cells and CD45^-^ TER119^-^CD31^-^ stromal cells, respectively (**Figure 2B-2C**). Phenotypically, skin Ebf2^+^ cells highly expressed PDGFRa/CD140a and SCA1 (PαS) whereas about 41% of the Ebf2^-^ cells were PαS cells (**Figure 2D**).

**Figure 2.**
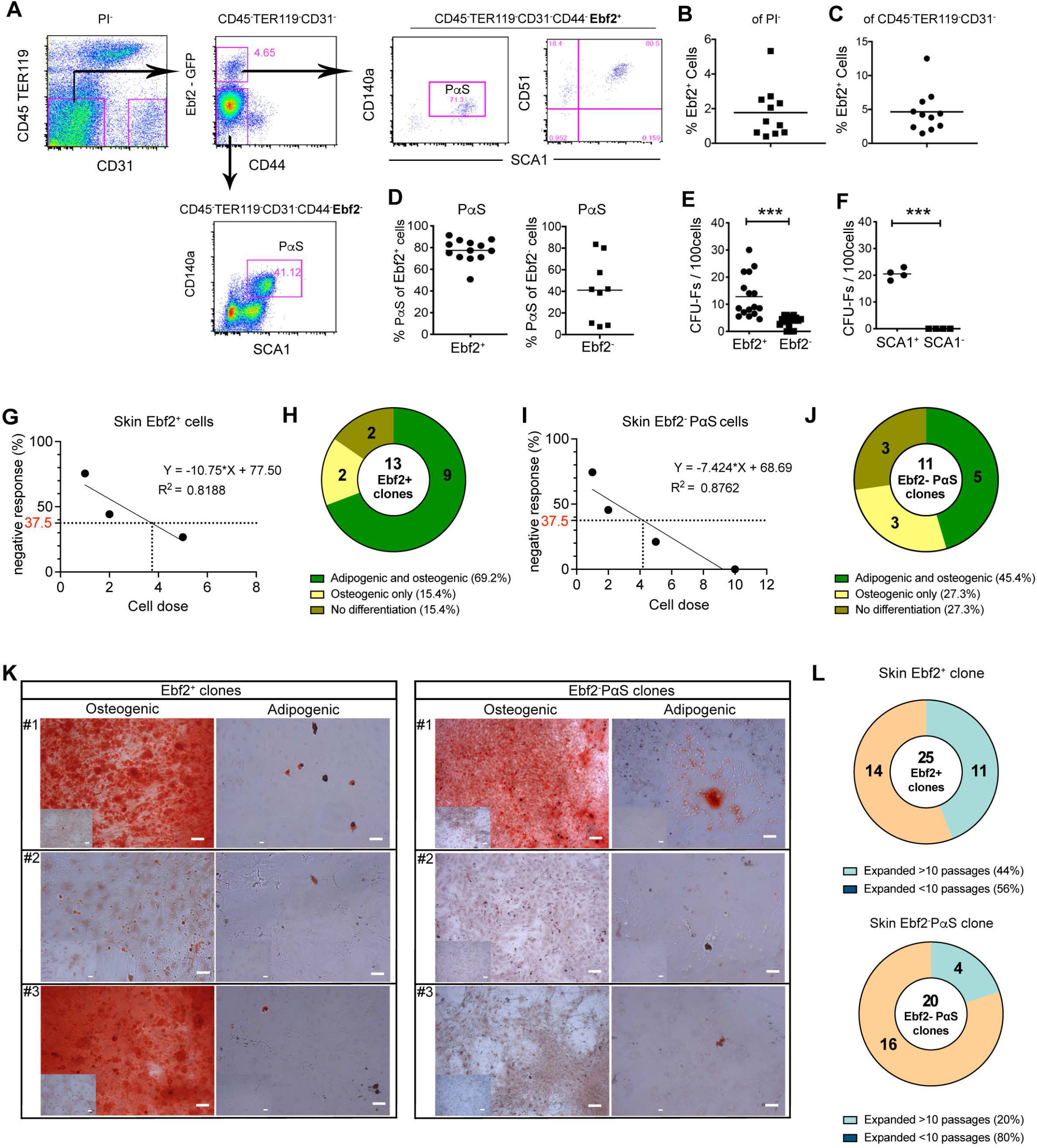
Identification of skin MPC subsets by Ebf2 expression. (**A**) A representative FACS profile showing FACS sorting/analysis of the Ebf2^+^ and Ebf2^-^ cells in dorsal skin. The cells were first gated within non-hematopoietic (CD45^-^TER119^-^) and non-endothelial (CD31^-^) live (PI^-^) stromal cells. These cells lacking expression of CD44 were further analyzed for their expression of SCA1, PDGFRa/CD140a (PαS) and CD51. The numbers in the panel are the mean frequencies. (**B-C**) The Ebf2^+^ cell frequency within total PI^-^ (**B**) or PI^-^CD45^-^TER119^-^CD31^-^ stromal cells (**C**) in dorsal skin. (**D**) The fractions of PαS cells within the Ebf2^+^ and Ebf2^-^ stromal cells. Each dot in **B-D** represents data from a single mouse in 3-6 experiments with the horizontal line as a mean value. (**E**) CFU-Fs in the Ebf2^+^ and Ebf2^-^ stromal cells. (**F**) CFU-Fs were exclusively found in the Ebf2^+^SCA1^+^ cell fraction. Data in **E-F** are from 3 independent experiments and each dot represents replicate assays from 2-3 mice in each experiment. The horizontal line represents mean value. *** P<0.001, **** P<0.0001, determined by Wilcoxon matched-signed pair rank test. **(G-J)** Single-cell analysis of CFU-Fs and lineage differentiation from the FACS-sorted Ebf2^+^ (**G-H**) and Ebf2^-^PαS (**I-J**) stromal cells. The CFU-F frequencies (**G**, **I**) were determined by limiting dilution at a density of 1, 2, 5, 10 cells per well in a 96-well plate and the frequency of the single cells with bi-lineage plasticity (**H, J**) were assessed by multilineage differentiation potential of single CFU-Fs derived from the cells. The simple linear regression was used to determine the dose responses in G and I. (**K**) Representative images of the osteogenic and adipogenic differentiation from single CFU- F clones derived from Ebf2^+^ and Ebf2^-^PαS cells. Scale bars in the images for Ebf2^+^ cells are 100 µm, and 250 µm and 100 µm in the images for osteogenic and adipogenic differentiation from Ebf2^-^PαS cells. (**L**) Population-doubling time (PDT) of randomly selected CFU-Fs derived from single Ebf2^+^ and Ebf2^-^PαS cells. Each line represents the growth kinetics of a single clone. See also in Figure S2.

Functionally, both the Ebf2^+^ and Ebf2^-^ cells contained CFU-Fs, a colony-forming unit characteristic for MSCs (**Figure 2E, Figure S2A**). The CFU-Fs in the Ebf2^+^ cells were generated exclusively by the SCA1^+^ population (**Figure 2F**). Both Ebf2^+^ and Ebf2^-^PαS cell populations showed similar proliferation kinetics after passage 5-6 (**Figure S2B**) and were able to differentiate towards osteogenic and adipogenic lineages *in vitro* albeit to a different degree (**Figure S2C-S2D**). However, they failed to differentiate into chondrocytes in either monolayer or micromass pellet culture in which condition BM MSCs could differentiate into chondrocytes (**Figure S2E**). These data indicate that the Ebf2^+^ and Ebf2^-^PαS cells in mouse skin may represent bipotential MPCs although they have the same immunophenotype as BM MSCs.

### Single-cell assay confirms osteo-adipo bipotentials of skin Ebf2^+^ and Ebf2^-^PαS cells

To validate the identity of the skin stromal subsets, we characterized these cells at single cell level. Limiting dilution experiment revealed that the CFU-F frequency in both skin Ebf2^+^ and Ebf2^-^PαS fractions was as high as 1 in 4 (**Figure 2G, 2I**). Of 13 randomly selected CFU-Fs from single Ebf2^+^ cells, 9 (69.2 %) showed both osteogenic and adipogenic differentiation potency although to different degrees (**Figure 2H, 2K**). However, only 45.4% (5 of 11) of the Ebf2^-^PαS cells showed bi-lineage potency toward osteoblasts and adipocytes (**Figure 2J-2K**). The chondrogenic potency, however, was not evaluated at single cell level since none of these cell subsets differentiated into chondrocytes at the bulk level. The self-renewal capacity of the Ebf2^+^ and Ebf2^-^ cells was evaluated by serially replating of single CFU-F derived from the single sorted cells. Eleven out of 25 CFU-Fs initially generated by single Ebf2^+^ cells could be serially replated for beyond 10 passages (**Figure 2L**). Meanwhile, only 4 in 20 CFU-F clones derived from Ebf2^-^PαS cells could be serially replated for over 10 passages (**Figure 2L**), indicating a limited self-renewal capacity of the Ebf2^-^PαS cells. These data further support the notion that the skin Ebf2^+^ and Ebf2^-^PαS cells are enriched with osteo-adipogenic MPCs.

### Skin Ebf2^+^ cells are mainly located at perivascular sites

Similar to that in mouse BM, the skin Ebf2^+^ cells did not express endothelial cell marker CD31 (**Figure S3A**). However, the Ebf2^+^ cells highly expressed PDGFRb/CD140b, a marker expressed in pericytes (Armulik et al., 2011) and niche-forming perivascular cells in BM (Kusumbe et al., 2016) (**Figure S3B**), pointing to a pericyte phenotype. Confocal imaging illustrated that the majority of the Ebf2^+^ cells were adjacent to the endothelial cells in the skin (**Figure 3A-3B**, **Figure S3C**). Some scattered Ebf2^+^ cells were also found in dermal panniculus carnosus, a layer of striated muscle (**Figure 3A**). Approximately 55% and 41% of the Ebf2^+^ cells were positive for the pericyte marker NG2, respectively (**Figure S3D, S3F**), and 81% of them expressed α-smooth muscle actin (SMA) (**Figure S3E-S3F**), reported to mark pericytes in BM (Baryawno et al., 2019). Altogether, these data suggest a perivascular origin and heterogeneity of the skin Ebf2^+^ cells.

**Figure 3.**
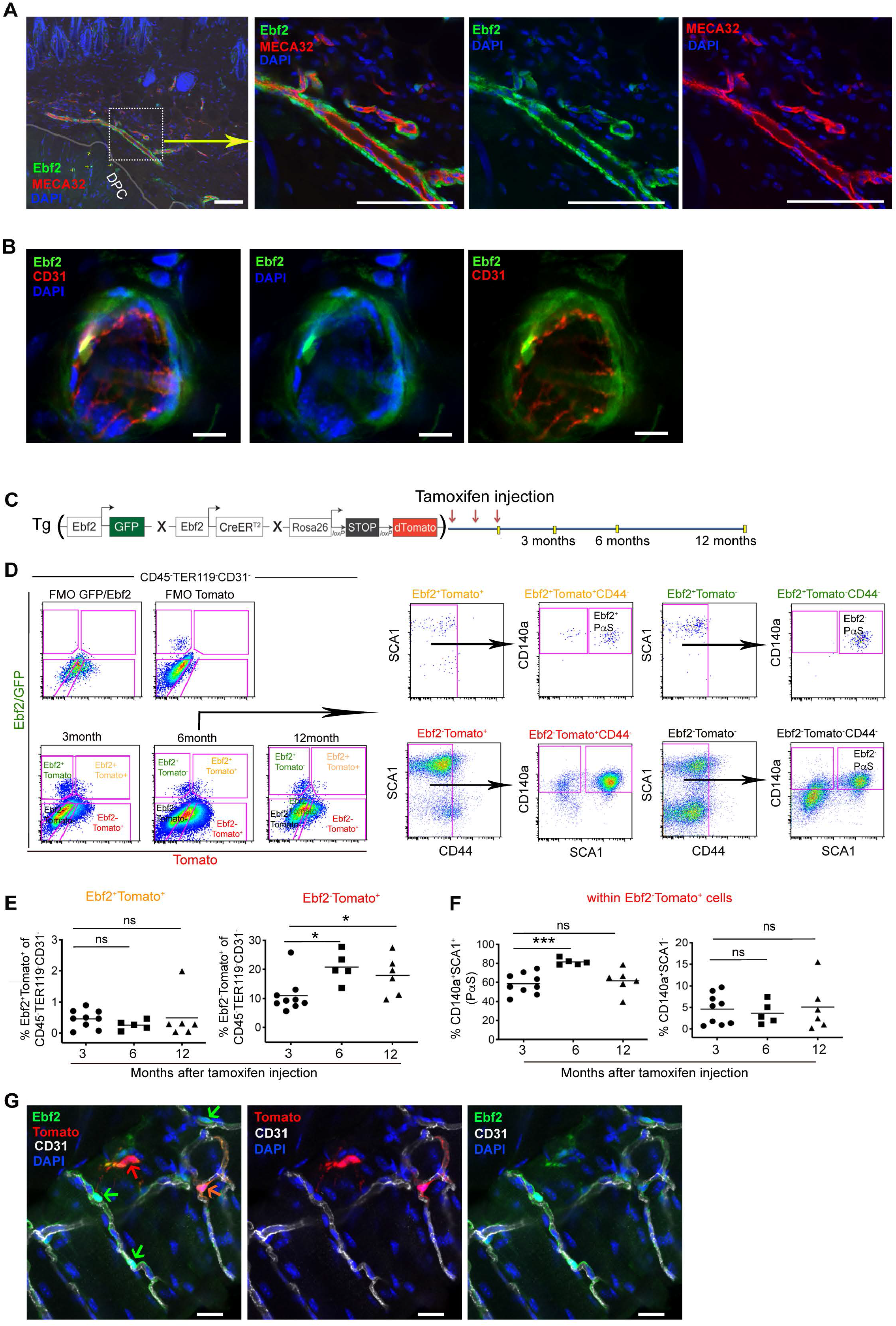
Distribution and hierarchical relationship of skin MPCs. (**A**) Localization of Ebf2^+^ cells in skin perivascular area and in dermal panniculus carnosus (DPC). The endothelial cells in the vessels were identified by MECA32 staining. Arrows point to the Ebf2^+^ cells in DPC area. Scale bars are 100 µm. (**B**) Representative image showing the perivascular localization of Ebf2^+^ cells adjacent to CD31^+^ endothelial cells in dorsal skin. Scale bars are 10 µm. (**C**) A scheme showing strategy for lineage-tracing of the Ebf2^+^ cells in skin. The Ebf2/GFP^+^Tomato^+^ cells and their progenies (Ebf2/GFP^-^Tomato^+^) were traced by FACS at 3, 6, 12 months after tamoxifen injection. (**D**) A representative FACS profile showing analysis of activated Ebf2^+^ cells (GFP^+^Tomato^+^) and their progeny (GFP^-^Tomato^+^). The gates for different cell subsets were defined with FMO from bi-transgenic or single transgenic mice. Each stromal cell subset (PI^-^CD45^-^TER119^-^ CD31^-^) was gated within the CD44^-^ fractions and subsequently gated for PαS cells based on CD140a and SCA1 expression. (**E**) The frequencies of Ebf2/GFP^+^Tomato^+^ cells and its progenies (Ebf2/GFP^-^Tomato^+^) within stromal cells. Each dot represents data from each mouse from 2-3 independent experiments. Horizontal bars represent the mean, * P<0.05, determined by unpaired *t*-test. (**F**) The fractions of PαS and Ebf2^-^CD140a^+^SCA1^-^ cells within total GFP^-^Tomato^+^ cells generated by Ebf2^+^ cells. Each dot represents data from each mouse from 2-3 independent experiments. Horizontal bars represent the mean, *** P<0.001, determined by unpaired *t*-test. (**G**) Distribution of the single Ebf2/GFP^+^Tomato^-^ cells (green), activated Ebf2/GFP^+^Tomato^+^ cells (orange) and Ebf2/GFP^-^Tomato^+^ (red) cells at 3 months after tamoxifen injection. Scale bars are 20 µm. See also in Figure S3-S4.

### The Ebf2^+^ cells in skin contribute to the mesenchymal cell turnover

To understand how mesenchymal cell niche in the skin is maintained, we determined the physiological contribution of the Ebf2^+^ stromal cells to the niche formation by lineage-tracing using Tg*Ebf2-Egfp xCre^ERT2^* X *Rosa26-tomato* mice (**Figure 3C**). This mouse model allowed us to track the fates of the Ebf2^+^ cells *in vivo* since the Ebf2^+^ cells express GFP and their progeny is marked by Tomato after tamoxifen (TAM) injection. About 15% of total Ebf2^+^ cells were Tomato^+^(Ebf2^+^Tomato^+^) cells at 3 months after TAM induction (**Figure S4A-S4B**). The Ebf2^+^Tomato^+^ cells could generate Ebf2^-^PαS MPCs and more mature CD45^-^TER119^-^CD31^-^ CD44^-^CD140a^+^SCA1^-^ stromal cells (**Figure 3D-3E**), indicating the important role of the Ebf2^+^ cells in maintaining the mesenchymal cell compartment in the skin.

The fraction of the Ebf2^+^Tomato^+^ cells within stromal cells remained constant up to 12 months post-TAM induction (**Figure 3E**), which is likely attributed to their self-renewal ability. However, the Ebf2^-^Tomato^+^ cells which were produced by the Ebf2^+^Tomato^+^ cells increased from 11% at 3 months to 21% at 6 months post-TAM injection, then remained stable (**Figure 3E**). Within the newly generated Ebf2^-^Tomato^+^ cells, the majority were PαS MPCs and about 5% were the CD140a^+^SCA1^-^ cells (**Figure 3F**). Inversely, within the total PαS MPCs, only 0.5% were Ebf2^+^Tomato^+^ cells and 30% were the newly generated Ebf2^-^Tomato^+^ PαS cells at 6 months post-TAM (**Figure S4C-S4E**). These data suggest a substantial contribution of the Ebf2^+^ MPCs to generating the Ebf2^-^Tomato^+^PαS MPCs while maintaining the Ebf2^+^ cell pool. Similar to the Ebf2^+^ cells, the Ebf2^-^Tomato^+^ cells were mainly distributed in the lower dermis and adjacent to CD31^+^ endothelial cells (**Figure 3G**). However, they lacked NG2 expression (**Figure S4F**). Only 19% and 28% of the Ebf2^-^Tomato^+^ cells expressed NESTIN and α-SMA, respectively (**Figure S4G-S4I**).

Altogether, skin harbors Ebf2^+^ and Ebf2^-^PαS MPCs with a similar phenotype as BM MSCs. The Ebf2^+^ MPCs reside at the top of the developmental hierarchy and contribute to mesenchymal cell generation in the skin.

### Skin MPCs play a role in maintaining normal hematopoietic and AML cells *in vitro*

Identification of the skin MPC populations made it possible to analyze their potential role in the maintenance of AML cells. We first evaluated the function of skin MPC subsets in supporting normal hematopoietic stem cells (HSCs) in a co-culture system and compared to BM MSCs (**Figure S5A**). Similar to their BM counterparts, both skin Ebf2^+^ and Ebf2^-^PαS MPCs were able to maintain HSC activities, indicated by similar numbers of CFU-Cs generated from the Lineage^-^ SCA1^+^KIT^+^(LSK)CD150^+^ cells after the co-culture with skin MPCs or BM MSCs (**Figure S5B-S5C**). Further, similar cobblestone area forming cells (CAFCs) were generated from the BM LSK cells co-cultured with the skin MPCs or BM MSCs (**Figure S5D- S5E**). The BM Ebf2^-^PαS cells displayed better capacity in supporting CAFC formation from the LSK cells than the BM Ebf2^+^ MSCs (**Figure S5F**). Altogether, these data indicated that skin MPCs possessed similar hematopoiesis supportive function as BM MSCs.

To assess the role of skin MPCs in supporting AML LSC growth, we next performed CAFC assay of MLL-AF9^+^ AML cells (**Figure 4A**). Like BM MSCs, skin MPC subsets could support CAFC formation from the AML cells, but to a lesser degree than the BM Ebf2^-^PαS cells (**Figure 4B-4C**).

**Figure 4.**
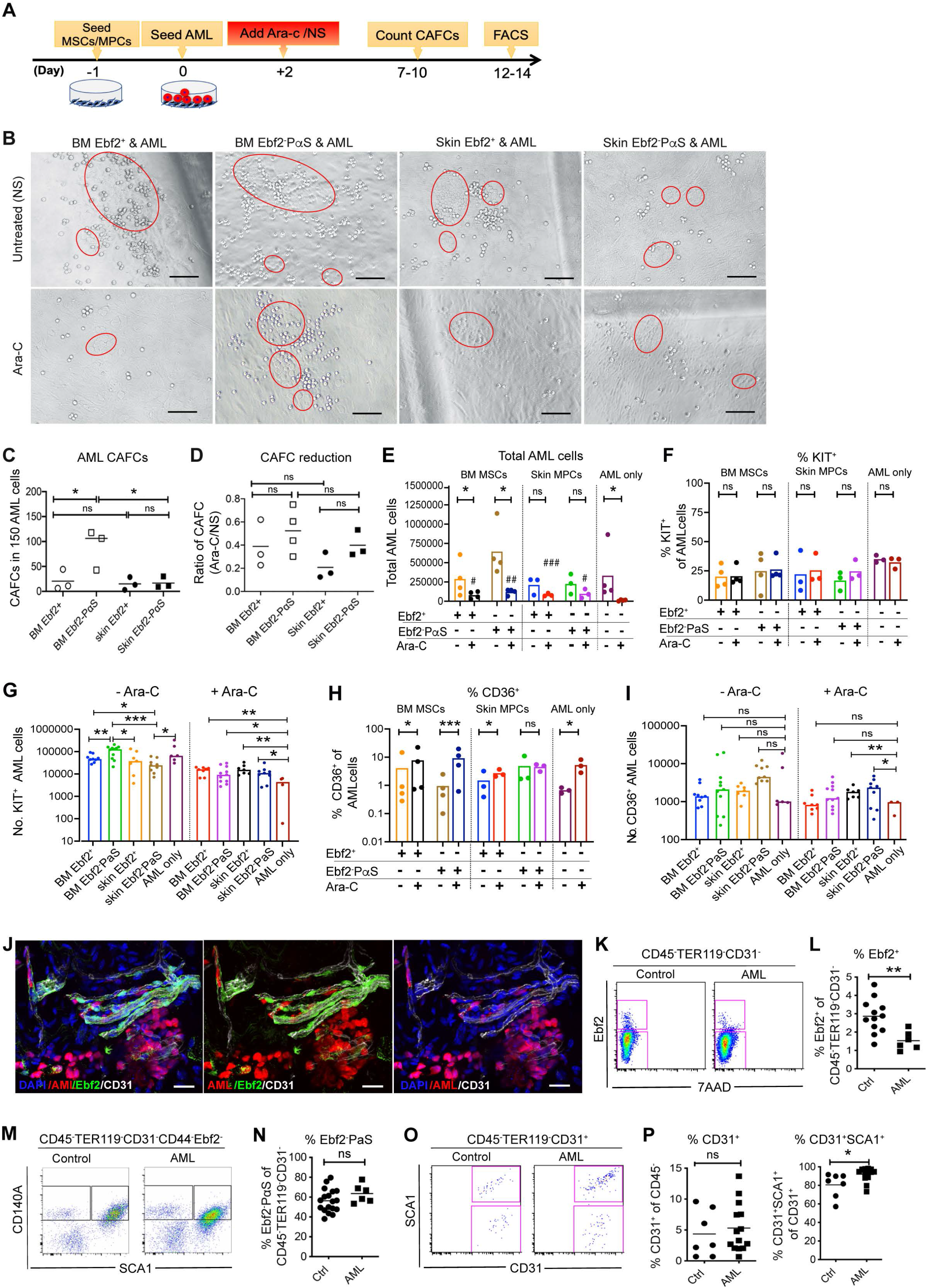
Skin MPCs support AML LSC growth and protect them from chemotherapy. (**A**) Experimental strategy for assessing the role of the skin MPC and BM MSC subsets for AML growth by cobblestone area-forming cell (CAFC) assay. (**B**) Representative images of CAFCs derived from the AML cells. Scale bars are 100 µm. (**C**) Total numbers of CAFCs generated from 150 MLL-AF9^+^ AML cells. (**D**) A proportion of residual CAFCs from AML cells after Ara-C treatment relative to the normal saline (NS)-treated controls. (**E**) The total number of the AML cells at 10 days after Ara-C treatment. (**F-G**) The frequency (F) and the numbers (G) of the KIT^+^ AML LSCs. (**H-I**) The frequency (H) and the numbers (I) of the CD36^+^ chemoresistant AML cells. Data in C-I were from 3-4 independent experiments and each dot represents the mean of replicate assays. The horizontal bars represent mean values. * P<0.05, ** P<0.01, *** P<0.001, determined by unpaired (C-D, G, I) or paired (E-F,H) *t*-test between Ara-C treated and nontreated co-cultures within the same stromal cell type. # P<0.05, ## P<0.01, ### P<0.001 determined by unpaired t test when comparing the Ara-C treated AML cells in co-cultures with BM MSCs or skin MPCs with the Ara-C treated AML monoculture. (**J**) Co-localization of AML cells (red) with the Ebf2^+^ cells. Ebf2 was determined by GFP (green) and the endothelial cells were marked by CD31 (white). Scale bars are 20 µm. (**K**) Representative FACS plot showing analysis of Ebf2^+^ MPCs in AML mouse dorsal skin. (**L**) The frequency of the Ebf2^+^ MPCs within stromal cells in dorsal skin tissue at endstage of AML. Data were from 6 independent experiments and each dot represents data from one mouse. The horizontal bars represent mean values. ** P<0.01, determined by unpaired *t*-test. (**M**) Representative FACS plot showing analysis of the skin Ebf2^-^PαS MPCs in healthy controls and AML mice. (**N**) The frequency of the Ebf2^-^PαS cells within stromal cells in dorsal skin at the end stage of AML. Data were from 6 independent experiments and each dot represents a mouse. The horizontal bars represent mean values. ns, no significant difference, determined by unpaired *t*- test. (**O**) FACS profile showing analysis of CD31^+^ cells in the dorsal skin of healthy and AML mice. (**P**) The frequency of total endothelial cells (CD31^+^) and arteriolar endothelial cells (CD31^+^SCA1^+^). ns, no significant difference, * P<0.05, determined by Mann-Whitney test.

Increasing evidence suggests that BM stromal cells may protect leukemia cells from cytostatic drugs (Alonso et al., 2015; Iwamoto et al., 2007; Jin et al., 2008; Xu et al., 2016; Zhang et al., 2013). However, it is not known if extramedullary stromal cells have similar functions. We then tested the chemoprotective function of skin MPCs by CAFC assay. We administered Ara-C 2 days after seeding the MLL-AF9^+^ AML cells to the MPCs to allow for efficient cell-cell interactions (**Figure 4A**). Both skin Ebf2^+^ and Ebf2^-^PαS cells could protect the AML LSCs from Ara-C treatment, demonstrated by the persistence of residual CAFCs after Ara-C treatment, in striking contrast to the efficient killing of the AML cells cultured alone (**Figure 4D-4E**). No difference in the frequency of KIT^+^ AML cells, representing AML LSCs (Somervaille and Cleary, 2006) was observed among all the co-cultures (**Figure 4F**). However, the numbers of the residual KIT^+^ LSCs were significantly increased in the co-cultures with the stromal cells compared to the monoculture post-Ara-C treatment (**Figure 4G**). CD36 was reported to be a marker for chemoresistant leukemic cells (Farge et al., 2017; Landberg et al., 2018; Ye et al., 2016). Post-Ara-C, the CD36^+^ AML cells were selectively enriched in all the co-cultures except that with skin Ebf2^-^PαS MPCs (**Figure 4H**), however, only in the culture with skin MPC subsets, significant more CD36^+^ AML cells were retained (**Figure 4I**). These data indicate a previously unrecognized role of skin MPCs in AML LSC maintenance and protection.

### The altered mesenchymal niches in the skin during AML development

Niche-remodeling in BM has been considered as one of the mechanisms contributing to leukemia progression (Ahsberg et al., 2020; Cai et al., 2022; Duarte et al., 2018; Xiao et al., 2018b). By confocal imaging, we observed the perivascular location of the AML cells (**Figure 1E**) and the localization adjacent to the skin Ebf2/GFP^+^ MPCs (**Figure 4J**). We then evaluated whether the skin cellular niches were remodeled by the AML cells. The proportion of Ebf2^+^ MPCs but not the Ebf2^-^PαS cells in the skin stromal cells were reduced post-symptomatic onset of AML (**Figure 4K-4N**). While this finding seems to be contrary to our previous observation of BM MSC increase in AML mouse BM, it is somewhat consistent with the reduced PαS cell fraction within the Ebf2^+^ cells in the AML mouse BM (Xiao et al., 2018b). In addition, while the total CD31^+^ endothelial cells were unaltered, as observed in BM (Xiao et al., 2018b), the CD31^+^SCA1^+^ arteriolar endothelial cells were significantly increased (**Figure 4O-4P**), indicating an increase of arterioles in dorsal skin during AML development.

### Molecular evidence for the role of skin MPCs in hematopoiesis and AML

To investigate the molecular mechanisms behind the hematopoiesis-supportive function of the skin MPCs, we performed RNA sequencing on the freshly sorted skin Ebf2^+^ and Ebf2^-^PαS MPCs. This revealed a distinct transcriptional profile of skin MPCs and BM MSCs with 1150 and 816 genes differentially expressed between skin Ebf2^+^ and Ebf2^-^PαS MPCs compared to BM MSCs, respectively (**Figure 5A**). However, only 135 genes were differentially expressed between the skin Ebf2^+^ and Ebf2^-^PαS cells. Among them, the genes associated with TNFα signaling, angiogenesis and epithelial to mesenchymal transition were upregulated while the cell proliferation genes were downregulated in the more primitive Ebf2^+^ cells (**Figure 5B**). We then compared skin MPC pool (Ebf2^+^ and Ebf2^-^PαS) with BM MSCs. The genes related to skin tissue maintenance (*Krt14*, *Krt19*) and adipogenesis (*Fabp4*) were upregulated in the skin MPCs (**Figure 5C-5D**), showing tissue origin-related characteristics of these cells. Consistent with the limited chondrogenic differentiation capacity of skin MPCs (**Figure S2C-S2D**), the genes associated with chondrogenesis were downregulated in the cells (**Figure 5E**). Conversely, skin MPCs were enriched with genes related to fatty acid metabolism and oxidative phosphorylation indicating a distinct metabolic profile between the skin and BM MSCs (**Figure 5D-5E**). Furthermore, the genes related to inflammatory response were enriched in the skin MPCs relative to BM MSCs (**Figure 5E-5F**), indicating a possible pro-inflammatory phenotype of skin MPCs.

**Figure 5.**
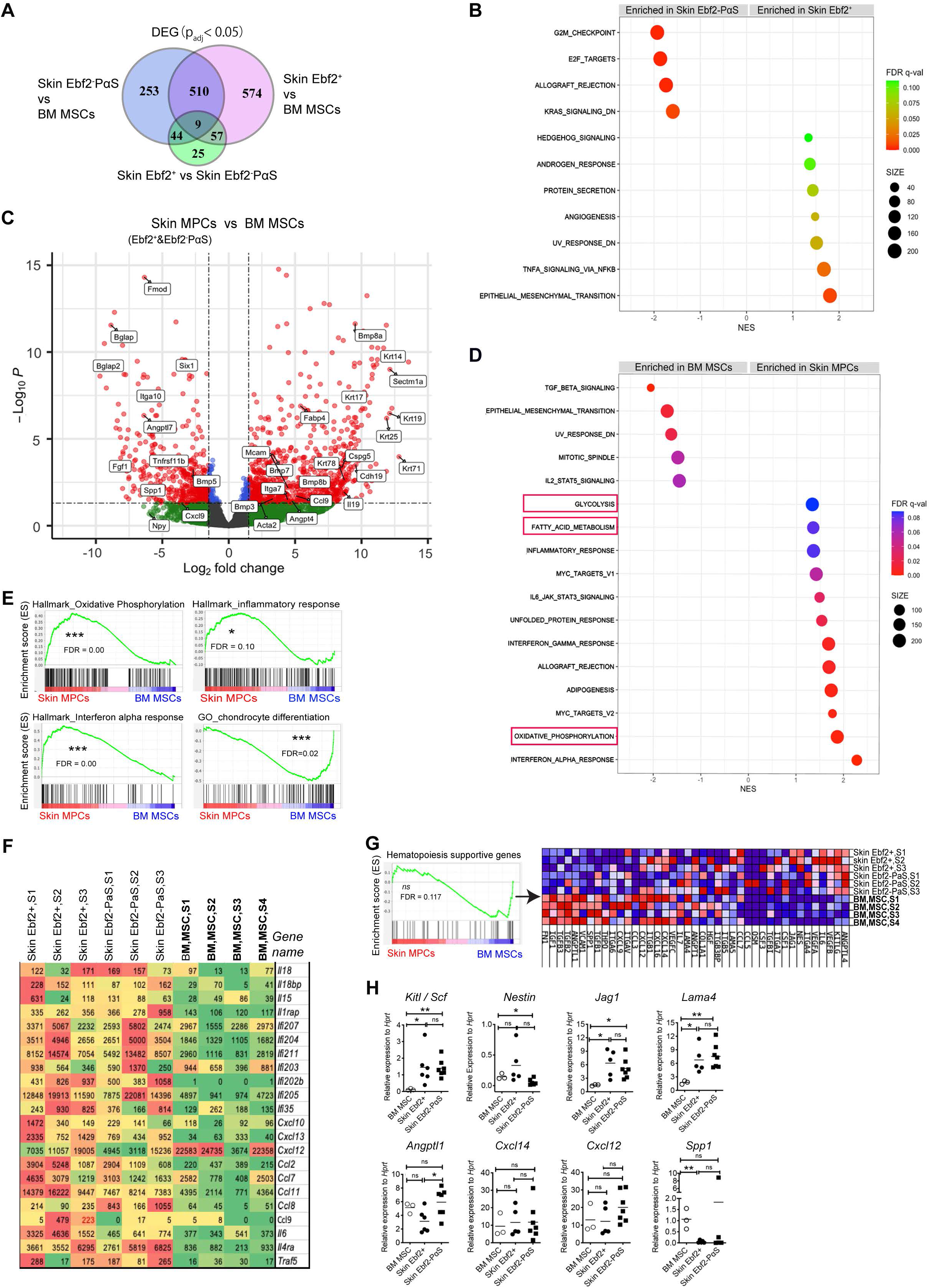
RNA sequencing revealed the molecular profile of skin Ebf2^+^ and Ebf2^-^PαS MPCs. (**A**) Venn diagram showing differentially expressed genes (DEG) among the skin Ebf2^+^ MPCs, skin Ebf2^-^PαS MPCs and BM MSCs. (**B**) Gene set enrichment analysis (GSEA) revealed the enrichment of genes associated with different biological processes and cellular responses in the skin Ebf2^+^ and Ebf2^-^PαS MPC subsets. (**C**) A volcano plot showing differentially expressed genes between skin MPCs (Ebf2^+^ and Ebf2^-^PαS) and BM MSCs. (**D**) GSEA revealed the enrichment of gene sets associated with various biological processes and cellular responses in the skin MPCs (Ebf2^+^ and Ebf2^-^PαS) and BM MSCs. (**E**) GSEA plots showing the enrichment of genes related to oxidative phosphorylation, inflammatory, interferon alpha response and chondrocyte differentiation in the skin MPCs compared to that in BM MSCs. FDR (false discovery rate). * P<0.05, *** P<0.001, determined by GSEA software. (**F**) Heatmap showing the expressions of selected inflammatory cytokines in skin Ebf2^+^ cells, Ebf2^-^PαS cells and BM MSCs. The heatmap was created in Excel using conditional formatting. The color scale was set based on the minimum, midpoint and maximum values of each gene in each row. Red correlates with high expression and green correlates with low expression. (**G**) Gene set enrichment plot showing hematopoiesis supportive niche genes in skin MPC subsets and BM MSCs, and the heatmap showing the gene expression levels. ns, no significant difference, determined by GSEA software. (**H**) Q-PCR of HSC niche genes in BM MSCs, skin Ebf2^+^ and Ebf2^-^PαS MPCs. Each dot represents mean of triplicate measurement of the gene expression relative to *Hprt*. Horizontal bars represent the mean values. Data were from 3 independent sorting experiments. ns, no significant difference, * P<0.05, ** P<0.01, determined by unpaired *t*-test.

Notably, there was no overall significant difference in the hematopoiesis supportive gene expression between the skin MPCs and BM MSCs (**Figure 5G**). The findings were confirmed by Q-PCR of the genes in these freshly sorted cells (**Figure 5H**). Similar to BM MSCs, both skin MPC subsets expressed *Cxcl12*, *Kitl*, *Jag1, Lama4* and *Angptl1,* all of which are known to be critical for hematopoiesis maintenance and regeneration (Arai et al., 2004; Calvi et al., 2003; Greenbaum et al., 2013; Gu et al., 2003; Omatsu et al., 2010; Poulos et al., 2013; Susek et al., 2018; Thoren et al., 2008). The expressions of *Kitl*, *Jag1* and *Lama4* were even significantly higher in skin Ebf2^+^ and Ebf2^-^PαS cells than that in BM MSCs. Moreover, the expression of *Spp1,* a negative regulator of HSC expansion (Stier et al., 2005) was lower in skin MPCs than that in BM MSCs (**Figure 5C, 5H**).These data provide molecular evidence for the supportive function of skin MPCs for HSCs and AML LSCs.

### Enhanced mitochondrial transfer from skin MPCs to AML cells associated with increased chemoprotection of AML cells

To reveal the mechanisms underlying the function of skin MPCs in maintaining and protecting AML LSCs, we examined the metabolic activity of skin MPCs in supporting AML cells since the genes related to oxidative phosphorylation and fatty acid metabolism are enriched in the skin MPCs. Mitochondria transfer has been reported as one of the protective mechanisms during chemotherapy in leukemias by providing energy support and anti-oxidant machinery to counteract oxidative stress induced by chemotherapy (Batsivari et al., 2020; Cai et al., 2022; Marlein et al., 2017; Moschoi et al., 2016). Consistent with their gene expression profile, our FACS analysis showed increased mitochondrial mass of skin MPCs compared to BM MSCs (**Figure 6A-6B**). Further, in a co-culture with mitochondria-prelabeled skin MPCs (**Figure 6C**), we detected enhanced mitochondrial transfer from skin MPCs to the AML cells compared to BM MSCs (**Figure 6D-6E**). Correspondingly, the AML cells co-cultured with skin MPCs showed lower ROS levels and higher viability during Ara-C treatment (**Figure 6F-6G**). The mitochondrial transfer and chemoprotection could be partially reversed by the treatment with microtubule inhibitor nocodazole and colchicine in the co-cultures (**Figure 6H-6J**). Similar to a previous report (Moschoi et al., 2016), the mitochondrial transfer seemed to be leukemia-specific since little or no mitochondrial transfer from skin MPCs to normal HSPCs (LSK BM cells) was detected in the co-cultures (**Figure 6K-6L**). Together, these data indicated that skin MPCs could maintain AML LSCs and protect them from Ara-C treatment *in vitro* partially by reducing oxidative stress.

**Figure 6.**
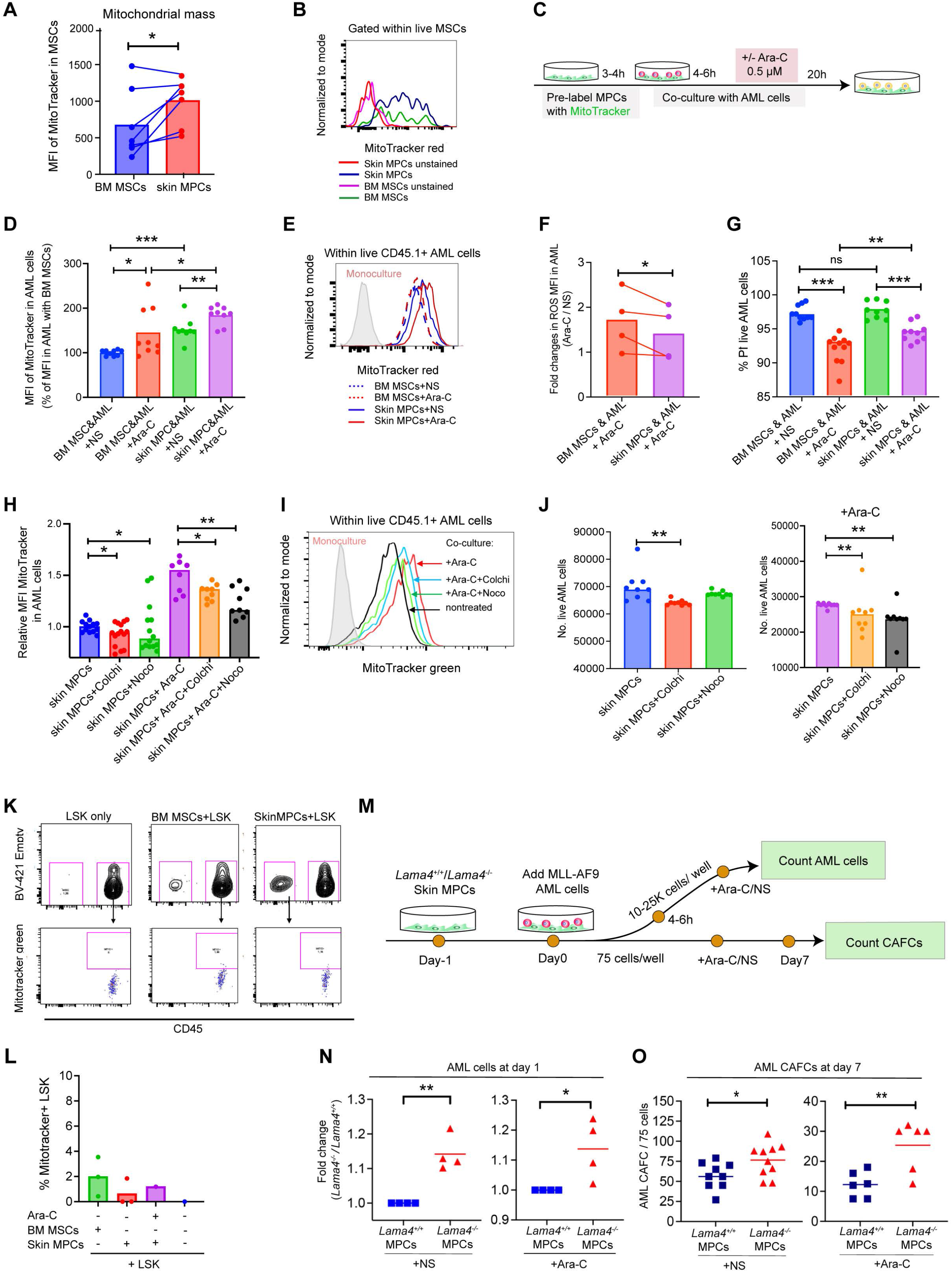
Mitochondrial transfer and Lama4 deficiency in skin MPCs contributed to the chemoprotection of AML cells from Ara-C. (**A-B**) Increased mitochondrial mass in skin MPCs compared to that in BM MSCs. (A) Mean fluorescence intensity (MFI) of MitoTracker red in BM MSCs and skin PαS MPCs. Each dot represents an individual assay in triplicate from 3 independent experiments. * P<0.05, determined by paired *t*-test. (B) Representative FACS histograms showing MitoTracker red staining in the skin MPCs and BM MSCs. (**C**) Experimental strategy for determining mitochondrial transfer from stromal cells to AML cells *in vitro*. Skin MPCs and BM MSCs that were pre-labelled with MitoTracker red were co- cultured with MLL-AF9^+^ AML cells for 24 hours. Ara-C was added 4-6 hours after seeding the AML cells. The stromal cell-derived mitochondria were detected by MFI of MitoTracker in the AML cells at 24h post-co-culture by FACS. (**D**) MFI of MitoTracker showing increased mitochondrial transfer from skin MPCs to AML cells than that from BM MSCs in the cocultures. Data shown are normalized MitoTracker MFI in the AML cells based on the values of that in the AML cells co-cultured with BM MSCs without Ara-C treatment. Each dot represents mean values of 3-4 replicated measurements in each experiment of 4. Horizontal bars are median values. * P<0.05, ** P<0.01, *** P<0.001, determined by Kolmogorov-Smirnov test. (**E**) Representative FACS histograms showing MitoTracker staining in the AML cells 24h post- co-culture with the stromal cells. The MFI from AML cells in the monoculture without the prelabeled MPCs was used as a negative control. (**F**) Reduced ROS levels in the AML cells co-cultured with skin PαS MPCs compared to that with BM MSCs 24h post-Ara-C treatment. Each dot represents the average value of triplicate assays from each experiment of 3. Horizontal bars are mean values. * P<0.05, determined by paired *t*-test. (**G**) The % of PI^-^ live AML cells in the cocultures 24h after Ara-C treatment. Each dot represents mean values of 3-4 replicated measurements in each experiment of 4. Horizontal bars are median values. The statistical differences were determined by unpaired *t*-test, ns, no significant difference, ** P<0.01, *** P<0.001, **** P<0.0001. (**H-J**) Impaired protective mitochondrial transfer from skin MPCs to AML cells in the cocultures after treatment with microtubule inhibitors. (**H**) MFIs of MitoTracker green in AML cells co-cultured with skin MPCs and treated with colchicine, nocodazole alone or in combination with Ara-C. The MFIs were normalized to the nontreated controls, from 4 independent experiments. Each dot represents a single measurement. *P<0.05, ** P<0.01, determined by unpaired *t-*test. (**I**) Representative histogram showing fluorescence intensity of mitotracker green in the AML cells. (**J**) The numbers of live AML cells co-cultured with skin MPCs after the treatments. ** P<0.01, determined by unpaired *t-*test. (**K-L**) Little mitochondrial transfer from skin or BM MSCs to normal BM HSPCs 24h post-coculture. The purified LIN^-^SCA1^+^KIT^+^ (LSK) cells were co-cultured with MitoTracker green prelabelled skin MPCs for 24h. The MitoTracker green^+^ LSK cells were analyzed post-co-culture based on CD45 expression, as shown in the FACS profile (K) and (L) the % of MitoTracker green^+^ LSK cells in co-cultures and monoculture. Each dot represents a replicate measurement. (**M**) Experimental layout for assessing the impact of *Lama4* loss in skin MPCs for AML growth *in vitro* using a co-culture system. The *Lama4*^+/+^ and *Lama4*^-/-^ skin MPCs were co-cultured with MLL-AF9 AML cells in a 96-well plate. For CAFC assay, Ara-C was added 2 days post-seeding of AML cells. The numbers of total AML cells and CAFCs were counted at 24-28h and day 7 post-seeding AML cells, respectively. (**N**) Fold changes in the number of the AML cells in the co-cultures with *Lama4*^-/-^ skin MPCs in relation to that with *Lama4*^+/+^ MPCs 24hours after Ara-C or NS treatment. Data were from 4 independent experiments and each dot represents the mean of triplicate assays. The horizontal bars represent mean values. *P<0.05, ** P<0.01, determined by paired *t*-test. (**O**) The number of CAFCs derived from AML cells in the co-cultures with *Lama4*^+/+^ and *Lama4*^-/-^ skin MPCs treated with NS or Ara-C. Data shown are triplicate values from 2-3 independent experiments. The horizontal bars represent mean values. *P<0.05, ** P<0.01, determined by unpaired *t*-test.

### *Lama4* loss in skin MPCs enhanced AML LSC proliferation and chemoprotection *in vitro*

*Lama4* was highly expressed in skin MPCs (**Figure 5H**) and we have found that *Lama4* loss in mice promoted AML LSC proliferation, chemoresistance, and relapse (Cai et al., 2022). This has tempted us to test the impact of *Lama4* deletion in skin MPCs on AML LSC growth (**Figure 6M**). We observed increased AML cell proliferation and chemoresistance to Ara-C at 24h post-co-culture with *Lama4^-/-^* skin MPCs (**Figure 6N**). Further CAFC assay indicated that *Lama4*^-/-^ skin MPCs promoted AML LSC proliferation and chemoresistance to Ara-C (**Figure 6O**), suggesting a suppressive impact of *Lama4* expression in skin MPCs on AML cell growth and chemoresistance.

### The increased chemoresistant AML cells in *Lama4*^-/-^ mouse skin post-Ara-C treatment

To further explore the molecular mechanism involving in the role of the extramedullary niche in the skin for maintaining residual AML cells after chemotherapy, we here took advantage of *Lama4*^-/-^ mice where AML progression and relapse are accelerated (Cai et al., 2022). To facilitate imaging of engrafted AML cells, DsRed-expressing MLL-AF9 AML cells were transplanted in sublethally irradiated *Lama4*^+/+^ and *Lama4*^-/-^ mice (**Figure 7A**). The mice were treated with either Ara-C or NS for 5 days at day 15 post-AML transplantation. Interestingly, one day post-Ara-C treatment, confocal imaging illustrated more AML cells in *Lama4*^-/-^ mouse skin, compared to that in *Lama4*^+/+^ mice following either NS or Ara-C treatment (**Figure 7B**). However, such a difference was not detected by FACS while there was a clear reduction of the AML cells in the PB and skin after Ara-C treatment (**Figure 7C**). It is possible that some of the remaining AML cells were lost during enzymatic digestion of the skin for FACS whereas confocal imaging was performed on the tissue fixed directly after dissecting. It is important to note that the chemoresistant CD36^+^ AML cells (Farge et al., 2017; Ye et al., 2016), not KIT^+^ cells were enriched in *Lama4*^-/-^ mouse skin post-Ara-C treatment (**Figure 7D**), which was not observed in the BM (Cai et al., 2022). It is worth mentioning that there was no homing preference of the AML cells to *Lama4*^-/-^ skin and BM compared to that in *Lama4*^+/+^ mice (**Figure 7E-7G**), suggesting that the observation of more AML cells in *Lama4*^-/-^ mouse skin was not related to the initial homing of the AML cells. Further, the AML cell homing to skin and BM was similar at 3 hours post-transplantation (**Figure 7H**), pointing towards a possibility that the skin tissue may act as an independent AML niche from the BM.

**Figure 7.**
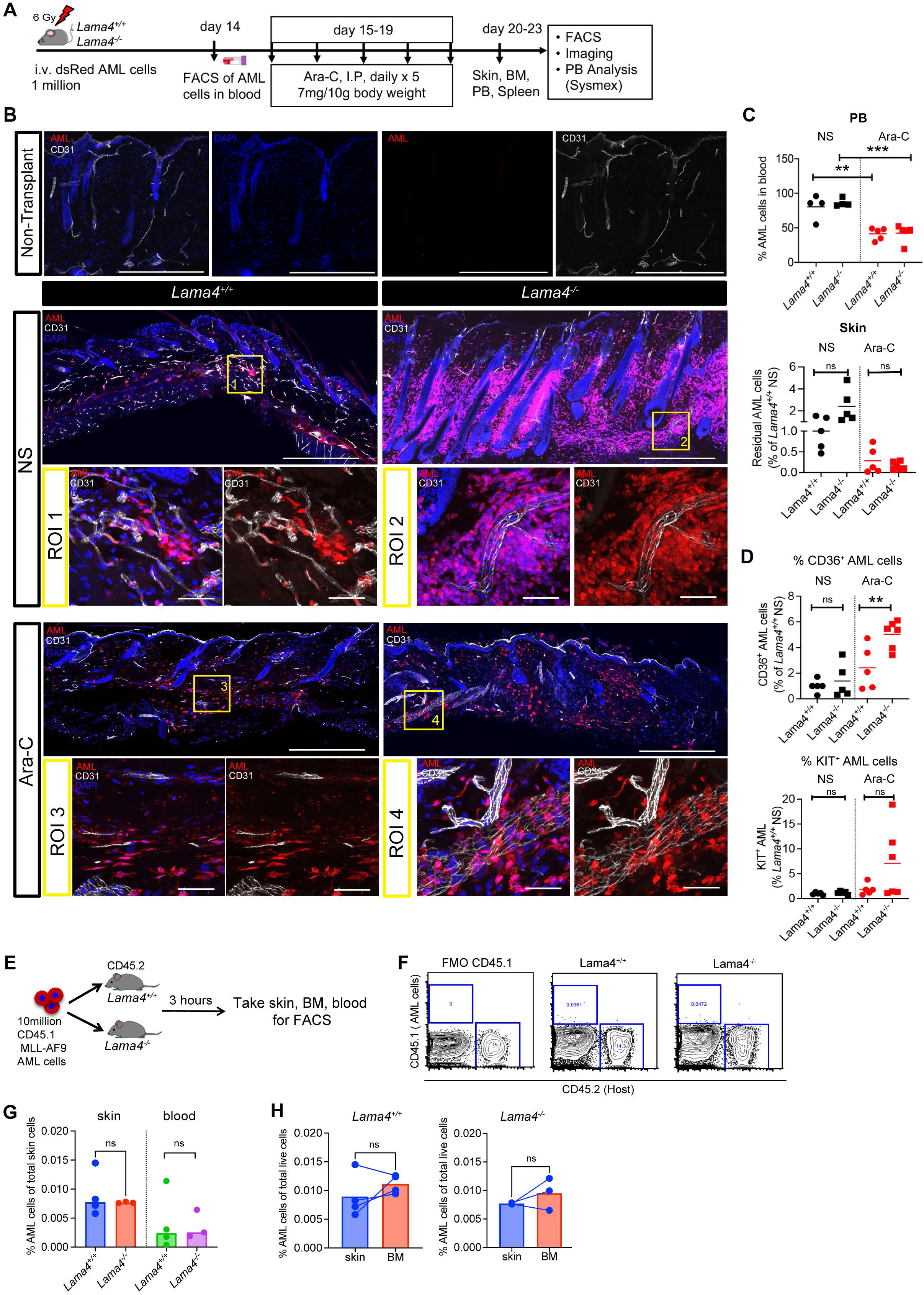
A high level of residual AML cell infiltration in skin tissue of AML-promoting *Lama4*^-/-^ mouse model post Ara-C treatment. (**A**) Experimental setup. *Lama4*^+/+^ and *Lama4*^-/-^ mice were first injected with DsRed-expressing MLL-AF9 AML cells after sublethal irradiation and treated with normal saline (NS) or Ara-C at day 15 after injection. Peripheral blood (PB) and BM were collected for FACS and confocal microscopy at 1-day after NS or Ara-C treatment. (**B**) Representative confocal images showing residual AML cells in dorsal skin at 1-day post- Ara-C treatment. Scale bars are 500 µm in the panoramic images and 50 µm in the region of interest (ROI) images (**C**) Frequency of AML cells in PB and skin at 1-day post-Ara-C treatment. (**D**) Proportion of CD36^+^ and KIT^+^ AML LSCs in skin at 1-day post-Ara-C treatment. Data were from 2-5 experiments and each dot in C-D represents data from a single mouse. The horizontal bars represent the mean values. n=5-6 per group. ** P<0.01, *** P<0.001, determined by unpaired *t*-test. (**E**) Experimental layout for assessing homing of the AML cells into *Lama4^+/+^*and *Lama4*^-/-^ mice 3 hours after transplantation. CD45.1^+^ MLL-AF9^+^ AML cells (10 million per mouse) were transplanted into the mice via tail vein injection without prior irradiation. Dorsal skin, bones, blood of the recipient mice were harvested 3 hours after AML cell transplantation and the homing of the CD45.1^+^ AML cells was examined by FACS based on CD45.1 expression. (**F**) Representative FACS profiles showing the gatings of CD45.1^+^ AML cells and CD45.2^+^ host cells. The gate for CD45.1^+^ cells was determined based on the FMO control without CD45.1 staining. (**G**) The frequencies of the AML cells in total live cells in the recipient skin and blood. ns, no significant difference, determined by unpaired *t*-test. The horizontal bars represent mean values. Each dot represents individual recipient mouse, n=3-4 per group. (**H**) The frequencies of the AML cells of total live cells in the *Lama4^+/+^* and *Lama4*^-/-^ recipient skin and BM. ns, no significant difference, determined by unpaired *t*-test. The horizontal bars represent mean values. Each dot represents individual recipient mouse.

To functionally evaluate AML-initiating LSCs, we performed serial transplantation with limited cell doses (10, 100, 1000 cells/mouse) and CFU-C assay post-Ara-C treatment (**Figure 8A**). Extreme limiting dilution analysis (Hu and Smyth, 2009) indicated no significant difference in the frequency of AML-initiating LSCs within the residual AML cells from *Lama4*^+/+^ and *Lama4*^-/-^ mouse skin (**Figure 8B-8C**). However, due to a relatively higher % of total AML cells in the donor *Lama4*^-/-^ mouse skin than that in the *Lama4*^+/+^ mice (**Figure 8D**), the calculated % of the AML-initiating LSCs and AML CFU-Cs in the *Lama4*^-/-^ skin appeared to be higher than that in the *Lama4*^+/+^ skin (**Figure 8E**). These data suggest that *Lama4* expression in skin mesenchymal niche plays a role in maintaining the chemosensitivity of AML cells in the skin to Ara-C.

**Figure 8.**
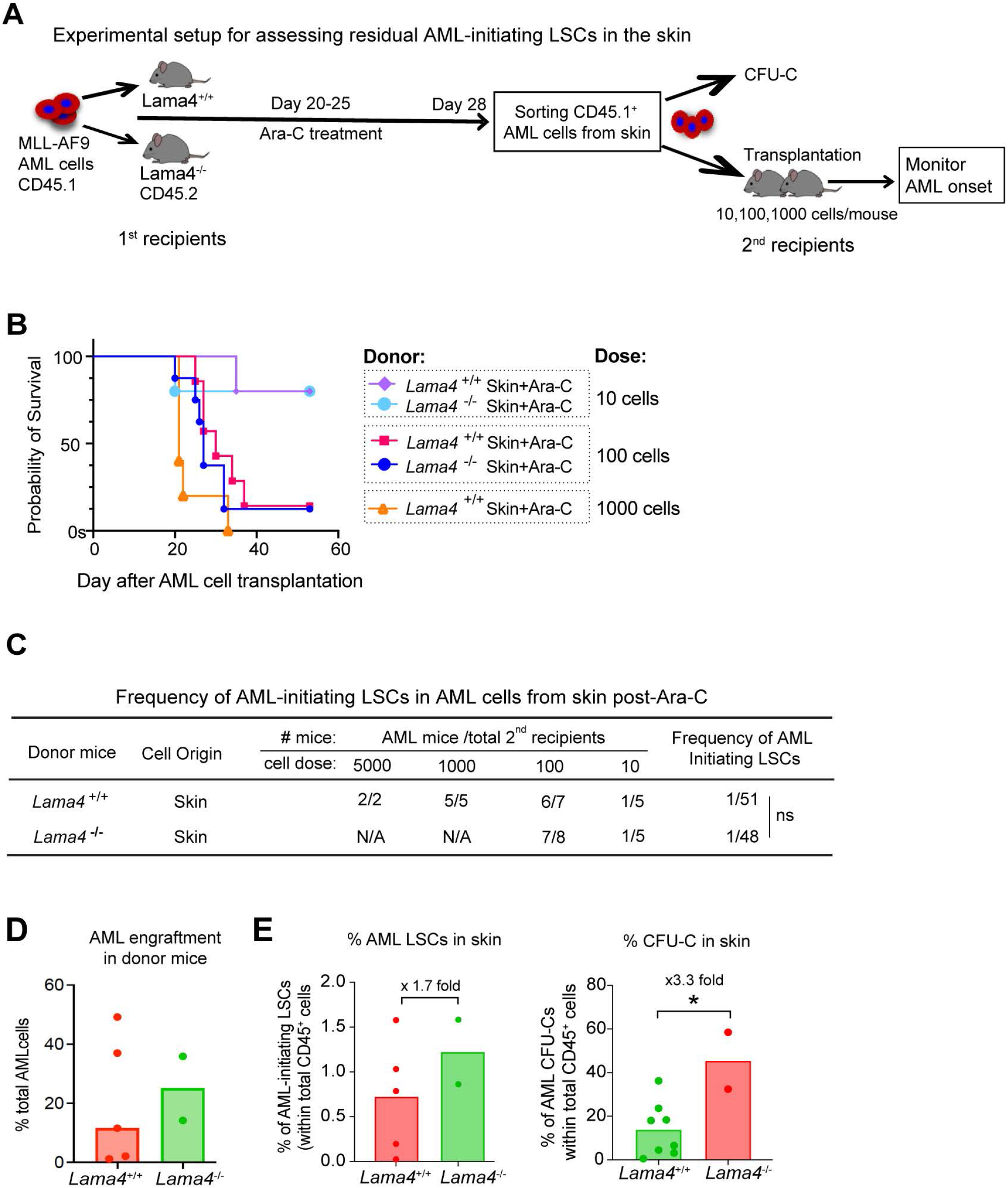
Serial transplantation indicated more residual AML-initiating LSCs in the *Lama4*^-/-^ mouse skin post-Ara-C treatment. (**A**) Experimental setup for determining the residual AML-initiating LSCs in the skin of *Lama4*^+/+^ and *Lama4*^-/-^ mice. The mice were injected with MLL-AF9 AML cells and treated with normal saline (NS) or Ara-C at day 20 after AML cell injection. The CD45.1^+^ AML cells were sorted from the skin at 3 days post-Ara-C treatment and intravenously transplanted in limiting cell doses (10, 100, 1000 cells/mouse) into secondary recipients without prior irradiation. The AML onset was monitored by FACS of AML cells in blood, spleen and BM. (**B**) The Kaplan-Meier survival curve of the secondary recipient mice which received skin-derived AML cells sorted from primary recipients after treatment with Ara-C. The dates were the days when the mice were found dead or moribund. The survival curve was generated by Log-rank (Mantel-Cox test). n=5-8 per group. (**C**) The frequencies of AML-initiating LSCs in the *Lama4*^+/+^ and *Lama4^-^*^/-^ mouse skin post- Ara-C treatment. The frequency was determined by Extreme Limiting dilution analysis based on the frequencies of the secondary mice that developed AML (Hu and Smyth, 2009). ns, no significant difference, determined by *Chi square* test. (**D**) The % of total AML cells in the skin of the donor *Lama4*^+/+^ and *Lama4^-^*^/-^ mice. Each dot represents data from a single mouse. The horizontal bars represent the median values. (**E**) The % of AML-initiating LSCs (left) and CFU-Cs (right) within total CD45^+^ cells in the skin. The data were calculated based on % of AML cells in skin and the frequency (**C**) of the LSCs and CFU-Cs, respectively. Each dot represents data from a single primary recipient mouse. The horizontal bars represent the mean values. * P<0.05, determined by unpaired *t*-test. Data were from 2 transplantation experiments.

## Discussion

Leukemia cutis is commonly observed in patients with monocytic AML and pediatric patients with congenital leukemia are prone to develop leukemia cutis (Parsi et al., 2020). Despite its association with a poor prognosis (Bakst et al., 2011; Gouache et al., 2018; Wang et al., 2019), the pathogenesis and impact of leukemia cutis on AML relapse remain largely unknown. We here have demonstrated in a transplantation-induced MLL-AF9 AML mouse model that the AML cells infiltrate in mouse skin, similar to leukemia cutis observed in patients with AML. Importantly, these skin-derived AML cells can efficiently regenerate AML post-transplantation even at a low dose (10 cells per mouse), indicating their high capacity in regenerating AML and pointing to a possible involvement of these leukemic cells in the relapse of AML.

Currently, little is known about the leukemia microenvironment or niche in the skin, which has limited our understanding of how leukemia cutis is maintained. We report that skin harbors two MPC subsets distinguished by Ebf2 expression, the Ebf2^+^ and Ebf2^-^PαS cells, sharing a similar phenotype with BM MSCs (Morikawa et al., 2009; Qian et al., 2013; Qian et al., 2012). The skin Ebf2^+^ MPCs co-express pericyte markers like CD140b/PDGFRb and α-SMA and show pericyte phenotype. The adjacent localization of the Ebf2^+^ cells to the AML cells in skin indicated a possibility that these cells act as a niche component for maintaining AML cells. This notion is supported by another important finding in this study that skin MPCs, similar to BM MSCs, show a supportive/protective function for AML LSCs. During AML progression, the skin mesenchymal cell niches are also altered with increased arteriolar endothelial cells and reduced Ebf2^+^, but not Ebf2^-^PαS MPCs after symptomatic AML onset. The niche-remodeling might in turn further promote AML progression since Ebf2^+^ cell deletion in mice accelerated AML development, as reported (Xiao et al., 2018b).

We have here attempted to address the potential mechanisms mediating the supportive role of skin MPCs. It is most likely that multiple mechanisms are involved in the AML protective role of skin MPCs. Mitochondrial transfer from stromal cells to leukemic cells has been reported to be one of the mechanisms to protect leukemic cells to survive from chemotherapy by providing an antioxidant system to counteract the therapy-induced oxidative stress (Forte et al., 2020). We find the increased mitochondrial mass in skin MPCs and enhanced mitochondrial transfer from skin MPCs to the AML cells during chemotherapy, compared to that in BM MSCs. Blocking mitochondrial transfer by microtubule inhibitors in the co-culture could compromise the protective effects of skin MPCs. These data suggest that skin MPCs may protect the AML cells by providing metabolic support via mitochondrial transfer. This notion was further supported by the enrichment of genes related to oxidative phosphorylation, fatty acid metabolism and glycolysis in skin MPCs. Furthermore, the skin MPCs express a higher level of *Lama4*, *Kitl* and *Jag1* than BM MSCs, which are known to be important for normal hematopoiesis maintenance and leukemic cell proliferation (Cai et al., 2022; Poulos et al., 2013; Somervaille and Cleary, 2006; Thoren et al., 2008). Our data indicate that *Lama4* expression in skin MPCs may have an impact on AML LSC chemosensitivity since *Lama4* deletion in skin MPCs led to enhanced proliferation and chemoprotection of AML LSCs. More work is required for assessing the potential impact of other niche factors like *Kitl, Jag1* and *Angptl*. In addition, the upregulation of inflammatory cytokines including interferon, *Il6*, *Il18*, *Il4* and *Cxcl10* in the skin MPCs points to another possible mechanism contributing to the AML maintenance in the skin, as these cytokines affect leukemic/cancer cell growth (Madapura et al., 2017; Manshouri et al., 2011; Pena-Martinez et al., 2018; Welner et al., 2015).

The origin of the AML cells in the skin remains an open question. In patients, leukemia cutis is usually present concurrently with AML cell infiltration in BM. However, in some cases, it may precede systemic involvement (Krooks and Weatherall, 2018). Therefore, it is unclear whether the leukemic cells infiltrated in the skin originate from BM. We here find that the AML engraftment in the skin does not correlate with that in the blood, suggesting AML cell maintenance in the skin may not be dependent on the circulating AML cells although they were initially derived from the injected AML cells. On the contrary, the AML burden in skin inversely correlates with the time of AML symptomatic onset, which is in line with the association of leukemia cutis with poor prognosis in patients with AML (Krooks and Weatherall, 2018).

In summary, we here report that skin harbors primitive Ebf2^+^ MPCs and the downstream Ebf2^-^ MPCs. These MPCs are colocalized with AML cells at perivascular sites during AML development. The residual AML cells in the skin after chemotherapy can regenerate leukemia post-transplantation, indicating the existence of AML LSCs in the skin. The skin MPCs can provide metabolic support for AML cells to protect them from chemotherapy *in vitro*. We for the first time provide evidence for the characteristics of skin mesenchymal cell niches and their roles in AML LSC growth and chemotherapy response, as well as the impact of *Lama4* expression in the niches on the chemoprotection of AML LSCs. These findings warrant future studies on the role of skin mesenchymal niches in chemotherapy response and relapse of AML patients.

## Methods

### Mice

*Ebf2-Egfp* reporter FVB/N mice (RRID:MGI:4421814) (Qian et al., 2013) at 8 to 14-week old were used. Transgenic *Ebf2-Egfp*X*Ebf2-Cre^ERT2^*X *Rosa26^loxp^Stop^loxp^-Tomato* mice were generated by crossing *Ebf2-Egfp* with *Ebf2-Cre^ERT2^*X*Rosa26^loxp^Stop^loxp^-Tomato* mice for lineage tracing. To activate Cre, tamoxifen (TAM) (Sigma) was intraperitoneally injected at 3 mg/20g body weight for 3 times every second day (Xiao et al., 2018b). *Lama4^-^*^/-^ C57BL/6 mice were generated as described (Thyboll et al., 2002). All mice were maintained in specific-pathogen-free conditions in the animal facility of Karolinska Institute. Animal procedures were approved by local ethics committee (ethical number 15861-2018) at Karolinska Institute (Stockholm, Sweden).

### Generation of MLL-AF9-induced AML mouse model

AML mouse model was induced as previously described (Xiao et al., 2018b). Briefly, 250.000 MLL-AF9 expressing cells were intravenously injected into non-irradiated mice to generate the AML mouse model. For detecting AML infiltration in dorsal skin tissue, 1×10^6^ of MLL-AF9 previously generated in DsRed C57BL/6J transgenic mice (Hartwell et al., 2013) were intravenously injected into sublethally irradiated (6 Gy) *Lama4^+^*^/+^ and *Lama4*^-/-^ mice, as described (Cai et al., 2022). At day 15, the mice were treated intraperitoneally with Cytarabine (Ara-C, Jena Bioscience) at the dose of 100mg/Kg or 700mg/kg body weight, daily for 5 consecutive days. For non-irradiated mice, the treatment started at day 20. The mice were sacrificed one day after the last injection for AML engraftment analysis in peripheral blood (PB), BM and dorsal skin by FACS. The distribution of residual AML cells in dorsal skin was visualized by confocal imaging.

### Skin MPC isolation by FACS

Mouse dorsal skin was minced and digested with 0.2% collagenase II (CLSII Worthington Biochemicals) in PBS supplemented with 20% FBS for 60 minutes at 37°C. After washing with PBS/20%FBS followed by PBS, the digested skin tissue was treated with 0.05% trypsin-EDTA (GIBCO) for 10 to 15 minutes at 37°C. Cell suspension was spun down at 300 g for 10 minutes. Mononuclear cells were then incubated with FcR (CD16/32) antibody and stained with CD45, TER119, CD31, CD44, CD140a, and SCA1 for 15 minutes at 2-8°C. Dead cells were excluded by propidium iodide (PI) staining. For sorting native skin MPCs, hematopoietic and endothelial populations were firstly excluded (CD45^-^TER119^-^CD31^-^) and Ebf2 gate was then defined based on Fluorescence Minus One (FMO) using skin mononuclear cells from a non-transgenic mouse. Skin Ebf2^+^ and Ebf2^-^ cells were sorted on a FACS Aria III Sorp (BD Biosciences, San Jose). See Table S1 for detailed antibody information.

### Serial transplantation

To test the leukemia-initiating capacity of the AML cells infiltrated in skin tissue, the AML CD45.1^+^ cells were sorted from skin tissue of the mice that have developed AML post-AML cell injection. The mice were treated with normal saline or Ara-C for 5 days at day 20-21 after AML cell injection. The AML cells in skin were collected 2-3 days after the last injection, sorted by FACS and transplanted into non-irradiated secondary C57BL/6J recipient mice via tail vein at a dose of 10, 100. 1000 or 5000 cells per mouse. The AML development was monitored by blood analysis using FACS and Sysmex as well as by assessing the general health conditions. The survival rate of the mice was estimated based on the date when the mice were found dead or in moribund status. Bones and blood were collected for determining AML engraftment.

### *In vivo* lineage tracing

This was performed as previously described (Xiao et al., 2018b). Dorsal skin from triple transgenic *Ebf2-Egfp* x *Ebf2-Cre^ERT2^*x *Rosa26^loxp^Stop^loxp^- Tomato* mice (Xiao et al., 2018b) were collected at 3, 6 and 12 months after tamoxifen injection. Approximately, 1 x 1 cm of dorsal skin tissue was fixed and embedded in OCT for whole mount immunofluorescence. For evaluating the frequency of Ebf2^+^ cells and their progeny (Tomato^+^), 2 x 2 cm of dorsal skin tissue was processed for skin MPC isolation by FACS (BD FACS Aria^TM^ III). For defining the gates for GFP^+^ and Tomato^+^ cells, cells from bi-transgenic or single transgenic mouse were used as FMO controls. After excluding dead cells by PI staining, the CD45^-^TER119^-^CD31^-^ stromal cells were gated. Subsequently, the CD44^-^ fractions were divided based on CD140a and SCA1 expression.

### Skin MPC expansion and proliferation kinetics *in vitro*

For *in vitro* expansion, CFU-Fs from FACS-sorted skin Ebf2^+^ and Ebf2^-^PαS cells were trypsinized with 0.05% trypsin-EDTA (GIBCO), collected, and plated to T-25 flasks at 400 cells /cm^2^. Cells were then expanded in complete Dulbecco modified Eagle Medium (DMEM, 31966, GIBCO) containing 10% FBS, 10 mM HEPES (1 M), 100 U of penicillin / streptomycin, and 10^-4^ M 2-Mercaptoethanol (M7522, Sigma) under hypoxic condition (1-2% O_2_) until 80%- 90% confluence was reached. Proliferation rates of skin Ebf2^+^ and Ebf2^-^PαS cells were evaluated by population doubling time (PDT) assays. PDT was calculated by dividing number of days for culturing the cells with number of population doublings (PDs) using following formula: PDT=Culture time (days)/PD where PD=log (NH/NI)/log2, NH is harvested cell number and NI is initial cell number.

### Dorsal skin wholemount immunofluorescence staining

Dorsal skin specimens were fixed in 4% PFA for 10 minutes followed by washing with PBS prior to embedding in OCT-Compound (Sakura Tissue Tex®, 4583). For immunofluorescence staining, 150 µm dorsal skin sections were first blocked with skim milk, fish skin gelatin (G7765, Sigma) and Triton X-100 (T8787, Sigma) for 1hour at RT. To visualize Ebf2^+^ cells in mouse dorsal skin, the sections were incubated with anti-GFP, additionally stained with either anti-mouse MECA32, CD31, NG2, NESTIN, or α-SMA antibodies. All stainings were performed at 2-8°C overnight prior to washing with PBS and staining with secondary antibody. Imaging was performed with confocal microscopy at Live Cell Imaging (LCI) facility after the section was mounted in Mountant PermaFlour (TA-030-F19, Thermo). See Table S2 for antibodies used for the confocal imaging.

Quantification of the Ebf2^+^ or Ebf2/GFP^-^Tomato^+^ cells expressing NESTIN, NG2 or α-SMA was performed with NIS-element AR-analysis ver 5.20.00 64-bit software (Nikon). From 100 µm section, 30-40 µm thickness was scanned with a confocal microscope to localize Ebf2^+^ and Ebf2^+^ progenies (Ebf2/GFP^-^Tomato^+^ cells). Areas where Ebf2^+^ and Ebf2/GFP^-^Tomato^+^ cells were distributed were further imaged every 1-1.1 µm depth to obtain Region of interest (ROI). Within ROI, positive expression of NESTIN, NG2 or α-SMA was defined based on control sections stained with secondary antibodies only while GFP^+^ signal was set based on samples from non-GFP reporter mice. Three to 4 mice were included for the quantification.

### RNA sequencing

Total RNA from freshly sorted skin Ebf2^+^ cells, skin Ebf2^-^ PαS cells and BM MSCs were isolated with RNeasy Micro Kit (Qiagen) according to the manufacturer’s protocol. cDNA was prepared using SMART-Seq v4 Ultra Low Input RNA Kit for Sequencing (634898, Takara Bio.). The cDNA quality was examined on Agilent TapeStation system (RRID:SCR_019547) using a High Sensitivity D5000 ScreenTape (5067-5592, Agilent). One ng cDNA was used for library preparation using *Nextera XT* DNA Library Preparation Kit (FC-131-1024 & FC-131- 1096, Illumina). The yield and quality of the amplified libraries were analyzed using Qubit by Thermo Fisher and the Agilent Tapestation System. The indexed cDNA libraries were sequenced on the Illumina 2000 or Nextseq 550 (Illumina, San Diego, CA) for a 75-cycle v2 sequencing run generating 75 bp single-end reads. About 7-20 million reads/sample were obtained. Sample quality was assessed using FastQC (v0.11.8) and MultiQC (v1.7). Reads were aligned to a reference built from Ensembl GRCm38 genome sequences using STAR (RRID:SCR_004463, v2.6.1d). All mapped counts to each gene were further calculated by FeatureCounts function from Subread package (Liao et al., 2013) installed in R. Genes with Reads Per Kilobase of transcript per Million mapped reads (RPKM) values more than 0.1 were considered as being actively transcribed and proceeded to the analysis of Differential Gene Expression (DGE) (Mortazavi et al., 2008). The normalized read counts assigned to each sample were generated by Deseq2 (RRID:SCR_015687). The differentially expressed genes between the cell subsets were identified by adjusted *P* value (*p*adj < 0.05) using Benjamini-Hochberg correction for multiple testing, together with thresholds at log_2_fold changes >1 (up- regulated) or <-1 (down-regulated). For the Gene Set Enrichment Analysis (GSEA), the normalized read counts were imported into the GSEA (v4.0.3) platform from Broad Institute (http://www.broadinstitute.org/gsea/index.jsp), with three gene sets being tested, including gene ontology (c5.all.v5.symbols.gmt), hallmark (h.all.v5.symbols.gmt) and KEGG (c2.kegg.v5.symbols.gmt). Gene sets tested were considered to be statistically enriched when the nominal *P* value < .01 and FDR < .25.

### Cobblestone area-forming cells (CAFC) assay using primary skin MPCs

This was done as previously described (Cai et al., 2022; van Os et al., 2008). Briefly, 30.000 skin/BM MPCs were first plated into a 24-multiwell plate and maintained in complete Dulbecco modified Eagle Medium (DMEM, 31966, GIBCO) containing 10% FBS, 10 mM HEPES (1M), 100U of penicillin/streptomycin, and 10^-4^M 2-Mercaptoethanol (M7522,Sigma) under hypoxic condition (1% O_2_). One day after, 150-300 MLL-AF9 or LSK cells were then added on the stromal cells in Myelocult (M5300, Stem Cell Technologies) with 10^-6^ M hydrocortisone (07904, Stem Cell Technologies), 1% penicillin streptomycin and 1ng/mL IL- 3 (R&D). The co-cultures were performed at 32°C in 5% CO_2_ for 7-14 days. A CAFC was defined as a cluster of more than 3 cells underneath the MSCs/MPCs. The CAFCs were visualized and scored with inverted microscope (CKX41, Olympus) on day 7 and 10. The cells were thereafter trypsinized and collected for FACS.

### Detection of reactive oxygen species (ROS) level and mitochondrial transfer

This was done as recently described (Cai et al., 2022). The MSCs/MPCs or AML cells were harvested and washed with PBS and incubated with 2 µM of H_2_-DCFDA (C6827, ThermoFisher) in DMEM at 37°C for 40 minutes to detect ROS level. The cells were then rinsed twice with PBS and resuspended with 150 µL of PI (1:1000) in 5%FBS/PBS. ROS levels were measured by FACS.

For mitochondrial transfer, MSCs or MPCs were stained with 100 nM MitoTracker™ red FM (M22426, ThermoFisher) at 37°C for 40 min, according to the manufacturer’s instructions. The cells were washed twice with PBS, then incubated for 3-4 hours to remove unbound probe before a final wash. Subsequently, 15,000-25,000 MLL-AF9 AML cells in Myelocult (M5300, StemCell Technologies) were plated and co-cultured with prelabelled MSCs for 24 hours. Mitochondrial transfer was quantified in AML cells (CD45.1^+^) by FACS and analyzed for MFI of the MitoTracker red.

### Multilineage differentiation assay

This was done as described (Xiao et al., 2018b). The skin Ebf2^+^, Ebf2^-^ and Ebf2^-^PαS cells were expanded in culture and plated at 400 cells/cm^2^ to each well of 24-well plate. For osteogenic differentiation, cells were cultured with complete osteogenic medium mixed by human/mouse StemXVivo osteogenic/adipogenic base medium (CCM007, R&D Systems) and mouse StemXVivo ostegenic supplement (CCM009; R&D Systems) under normoxic condition for 14-21 days. Differentiation toward osteoblast was evaluated by 1% alizarin red S (A5533, Sigma) staining after cell fixation with cold methanol. For adipogenic differentiation, the culture was performed with DMEM containing 10% FBS, 10 mM HEPES (1M), 100U of penicillin/streptomycin, 10^-4^ M 2-Mercaptoethanol (Sigma, M7522), 5 μg/mL insulin (I6634, Sigma), 20 μM Indomethacin (I7378, Sigma), 0.0115 mg/mL isobutylmethylxanthine (I-7018, Sigma), and 10^-6^ M dexamethasone (D2915, Sigma) for 2-3 weeks. Cells were first stained with 500 μL of 2 μM of Bodipy™ 500/510 (Invitrogen, D3823) for imaging live adipocytes, then fixed with 10% formalin and stained with 0.5% Oil Red O (O1391, Sigma). Chondrogenic differentiation was induced in monolayer culture where cells were cultured in 37°C under hypoxic condition (2% O_2_) with DMEM high glucose containing 10 mM HEPES (1M), 100U of penicillin/streptomycin, 10^-4^ M 2-Mercaptoethanol (M7522, Sigma), 2 mM pyruvate (P5280, Sigma), 0.35 mM L-proline (P5607-25G, Sigma), ITS^+3^ (I-3146, Sigma), 5 μg/mL L-ascorbic acid 2-phosphate (A7506, Sigma), 10^-7^ M dexamethasone (D2915, Sigma), and 10 ng/mL TGF-β3 (100-36E, Peprotech). To assess chondrogenic differentiation, toluidine blue (T3260, Sigma) (pH 2.0 to 2.5) was used to stain proteoglycan. Images were then taken under inverted microscope (CKX41, Olympus).

### *In vitro* chondrogenic induction in micromass pellet

After *in vitro* expansion, 2.5×10^5^ of BM MSCs or skin Ebf2^+^ or skin Ebf2^-^PαS MPCs were collected in 15mL tube and spun down at 300 g for 7 minutes. After removal of supernatant, 500 μM of chondrogenic medium consisting of DMEM high glucose with 10 mM HEPES, 100U of penicillin/streptomycin, 10^-4^ M 2-Mercaptoethanol (M7522, Sigma), 2 mM pyruvate (P5280, Sigma), 0.35 mM L-proline (P5607-25G, Sigma), ITS^+3^ (I-3146, Sigma), 5 μg/mL L-ascorbic acid 2-phosphate (A7506, Sigma), 10^-7^ M dexamethasone (D2915, Sigma,), and 10 ng/mL TGF-β3 (100-36E, Peprotech) were added. The chondrogenic induction was performed in 37°C under hypoxic condition (2% O_2_). Every 2-3 days, the medium was replaced until day 28. For evaluating the chondrogenic induction, the micromass pellets were washed with PBS prior to being fixed in 4% PFA for 2 days. After dehydration with 70% ethanol, the pellets were stained with toluidine blue (T3260, Sigma) at pH 2.0 to 2.5 for 15–30 minutes. Thereafter, the stained pellets were washed with 70% ethanol and embedded in OCT-compound (4583, Sakura Tissue Tex®). To visualize the formation of proteoglycan, the embedded pellet was cut to 10μm, mounted and observed by an inverted microscope (Axio Observer.Z1, Zeis). Images were processed with Zen software (Carl Zeiss Microscopy, GmbH 2011).

### LIN^-^SCA1^+^KIT^+^ (LSK) CD150^+^ cell isolation

The LSKCD150^+^ HSCs were isolated from femurs and tibias of young adult mice, as described (Xiao et al., 2018a). See Table S1 for antibody detail information.

### Co-culture of skin MPCs with LSKCD150^+^ cells and colony-forming unit in culture (CFU-C) assay

Twenty-five thousand (25,000) expanded Ebf2^+^ or Ebf2^-^PαS MPCs from skin and BM at passage 4-13 were plated into each well of a 12-well plate and maintained in complete Dulbecco modified Eagle Medium (DMEM, 31966, GIBCO) containing 10% FBS, 10 mM HEPES (1M), 100U of penicillin/streptomycin and 10^-4^ M 2-Mercaptoethanol (M7522, Sigma) under hypoxic condition (2% O_2_) for 24h prior co-culture. The co-cultures were started by plating 2000 sorted LSKCD150^+^ to each well in Myelocult medium (M5300, Stem Cell Technologies) supplemented with 10^-6^ M hydrocortisone (74142, Stem Cell Technologies) and 1% penicillin streptomycin. After a 3-day co-culture in 5% CO_2_ at 37°C, the medium containing hematopoietic cells and trypsinized cells were collected for FACS analysis and CFU-C assay in Methylcellulose M3434 (Stem Cell Technologies). After 10 days of culture at 37°C in 5% CO_2,_ the CFU-Cs were stained with 2,7-diaminofluorene (DAF staining) and scored with an inverted microscope (CKX41, Olympus). A cell cluster with a minimum of 50 cells is defined as one colony. Colonies with DAF staining were defined as erythrocytes-containing colonies including CFU-GME and BFU-E, as performed (McGuckin et al., 2003; Xiao et al., 2018a).

### Statistical analysis

Unless mentioned, either unpaired t-test or Mann-Whitney test was used to determine the difference between the groups or cell subsets. All statistical tests were performed with GraphPad Prism 8 software (RRID: SCR_002798) with P<0.05 considered statistically significant.

### Online supplemental material

Fig. S1 shows AML cell distribution, absolute numbers in skin, BM, spleen and blood from AML mice under steady state and post-chemotherapy. Fig. S2 illustrates *in vitro* expansion and differentiation of the Skin Ebf2^+^ and Ebf2^-^ PαS cell subsets. Fig. S3 provides additional evidence for the perivascular location and expression of α-SMA and NG2 of skin Ebf2^+^ cells. Fig. S4 shows additional data from lineage-tracing studies suggesting that skin Ebf2^+^ cells generated Ebf2^-^ cells while maintaining themselves *in vivo*. Fig. S5 shows the data from in vitro co-culture experiments indicating that skin MPCs have a similar hematopoiesis supportive function to BM MSCs.

## Data availability

The RNA sequencing data in Fig. 5 are available at GEO under accession number GSE167562. All other data are available in the article or upon reasonable request to the corresponding author.

## Acknowledgements

We thank Tina Jacob, Karl Annusver and Alexandra Are at Department of Cell and Molecular Biology, Karolinska Institute, Sweden for valuable advice and technical discussion. We thank Alexandre Piccini, Yuanyuan Zhang and Andranik Durguryan in our group for technical help for the cell culture. We also thank Prof. Marja Ekblom, Lund University for her scientific input. This study was supported by the Swedish Research Council (2019-01361,2022-01228), Swedish Childhood Cancer Society (PR2015-0142, FoAss13/015, PR2017/0154, PR2020- 0166), Swedish Cancer Society (CAN2017/774, 19 0092 SIA and 20 1222PjF), Åke Olsson foundation, Radiumhemmets Forskningsfonder (151241, 171162, 191253) and Karolinska Institute Wallenberg Institute for Regenerative Medicine, Karolinska Institute doctoral education (KID) funding (2-1293/2014, 2021-00480), Stiftelsen Clas Groschinskys Minnesfond (ref:M16 50), Cancer Research KI (Karolinska Institute) and Incyte Biosciences Nordic to H.Q., and the Swedish Research Council (2018-02963) and Swedish Cancer Society (CAN 2018/793) to MK (Kasper). FACS analysis and cell sorting were performed at MedH Flow Cytometry core facility (Karolinska Institute), supported by KI/SLL. The confocal images were obtained at the LCI facility/Nikon Center of Excellence, Karolinska Institute, supported by grants from the Knut and Alice Wallenberg Foundation, the Swedish Research Council, KI infrastructure, Centre for Innovative Medicine and Jonasson center at the Royal Institute of Technology. All major computations were performed on resources provided by the Swedish National Infrastructure for Computing (SNIC) through Uppsala Multidisciplinary Center for Advanced Computational Science (UPPMAX) under Project sens2018540. We also would like to thank the core facility at NEO, BEA, Bioinformatics and Expression Analysis, which is supported by the board of research at the Karolinska Institute and the research committee at the Karolinska Hospital.

## Author contributions

LS has substantially participated in designing, performing experiments, collecting and analyzing data and manuscript writing. HC, EL, PX, MK(Kondo), XS, LT, AM and ASJ performed experiments, collected data and assisted with data analysis and manuscript-editing. MK (Kasper) provided scientific input and assistance with confocal imaging of skin tissues. MJ provided the DsRed-expressing MLL-AF9 AML cells and scientific input on the manuscript. KT provided Lama4^-/-^ mouse models and scientific input. HQ designed, performed experiments, collected and analyzed data, wrote the manuscript. All authors have approved the final version of the manuscript.

## Conflict of interest

All authors declare no competing financial interest regarding this study.

## Non-standard abbreviations

AML: acute myeloid leukemia
MSCs: mesenchymal stem cells
MPCs: mesenchymal progenitor cells
LSCs: leukemia-initiating stem cells
HSCs: hematopoietic stem cells
HSPCs: hematopoietic stem and progenitor cells
Ebf2: Early B-cell factor 2
BM: bone marrow
Ara-C: cytarabine
NS: normal saline
FACS: fluorescence- activated cell sorting
CFU-C: colony forming unit in culture
CAFC: cobblestone area-forming cells
DPC: dermal panniculus carnosus
CFU-F: colony-forming unit in fibroblast
PαS: PDGFRa/CD140a and SCA1-expressing stromal cells.

## Notes

https://www.ncbi.nlm.nih.gov/geo/query/acc.cgi?acc=GSE167562

### Competing Interest Statement

The authors have declared no competing interest.

### Summary of Updates

The title, Figure 1, Figure 6 and Figure 7 of this paper are updated. Figure 8 is added and supplemental figures are reduced to 5 by incorporating some of the supplemental figures into the main Figure 5 and Figure 7. The abstract, results, and discussion are updated, accordingly. This paper is now accepted by the Journal of Experimental Medicine.

