## Supplemental Figures and tables for "Skin mesenchymal niches maintain and protect AML-initiating stem cells"

Running title: Extramedullary mesenchymal cell niche in AML

Lakshmi Sandhow[1], Huan Cai[1#], Elory Leonard[1#], Pingnan Xiao[1], Luana Tomaipitinca[1], Alma Månsson[1], Makoto Kondo[1&], Xiaoyan Sun[2], Anne-Sofie Johansson[1], Karl Tryggvason[3], Maria Kasper[2], Marcus Järås[4], Hong Qian[1*]

Supplemental Figures 1-5

**Supplementary Figure 1 AML cell distribution in skin, BM, spleen and blood from AML mice under steady state and post-chemotherapy.** Data were from 2 independent experiments and each dot represents a single mouse with AML. The mice were treated with cytarabine (Ara-C) at day 20 post-AML transplantation for 5 days and the AML engraftment was analyzed at 3 days after the treatment.

**(A)** Hematoxylin and eosin (HE) staining of skin derived from a healthy and AML mouse. Scale bars are 50 µm.

**(B)** No significant correlation between AML engraftment in the skin and that in blood. No significant difference (ns) was determined by Spearman correlation.

(**C**) Total cellularity in the skin of mice with AML and healthy controls (Ctrl). Each dot represents data from one mouse. ns, no significant difference was determined by unpaired *t* test.

**(D)** The absolute numbers of AML cells in blood, skin, BM and spleen.

**(E)** The frequency of CFU-Cs derived from the AML cells in BM, skin and spleen of nontreated (left) and Ara-C-treated (right) mice. The frequencies were calculated based on the frequencies of CFU-Cs within the sorted AML cells and the % AML cells in each tissue. Each dot represents data from one mouse. * $P<0.05$, ** $P<0.01$, *** $P<0.0001$, determined by unpaired *t*-test (B, D, left) and paired *t*-test (D, right).

**Supplementary Figure 2 *In vitro* expansion and differentiation of the Skin Ebf2$^{+}$ and Ebf2$^{-}$ PαS cell subsets.**

(**A**) Images of CFU-Fs derived from the Ebf2$^{+}$ and Ebf2$^{-}$ cell subsets. Scale bars are 500 µm.

(**B**) Population doubling time (PDT) of the skin CD45$^{-}$TER119$^{-}$CD31$^{-}$Ebf2$^{+}$ and Ebf2$^{-}$ stromal cells in culture. PDT was calculated based on culture time (CT)/Cell doubling (CD) where CD=log (NH/NI)/ log 2, NH is harvested cell number and NI is initial cell number.

(**C**) Representative images of osteogenic, adipogenic and chondrogenic differentiation of the skin Ebf2$^{+}$ and Ebf2$^{-}$PαS cells. Scale bars are 500 µm (osteogenic), 100 µm (adipogenic) and 20 µm (chondrogenic).

(**D**) Bodipy™ 500/510 and oil red O staining of differentiated adipocytes from the skin Ebf2$^{+}$ and Ebf2$^{-}$PαS cells 21 days post the induction. Green and red represent adipocytes stained with bodipy 500/510 and oil red O, respectively. Scale bars are 100 µm.

(**E**) Representative images of Toluidine blue staining on micromass pellet after chondrogenic induction *in vitro.* The chondrogenic differentiation in a micromass pellet culture were performed on culture expanded BM MSCs and skin Ebf2$^{+}$ and Ebf2$^{-}$ cell subsets. Scale bars are 20 µm.

Related to Figure 2.

**Supplementary Figure 3 Skin Ebf2$^+$ cells are perivascular cells.**

(**A**) Frequencies of Ebf2$^+$ cells expressing CD31. Data are from 5 sorting experiments. Each dot represents one skin sample from one mouse.

(**B**) Representative FACS plots showing analysis of CD140b expression in the skin Ebf2$^+$ and Ebf2$^-$PαS cells.

(**C**) Perivascular localization of Ebf2$^+$/GFP$^+$ cells. The vessels were identified by MECA32 staining in mouse dorsal skin tissue. Scale bars are 20 μm.

**(D-E**) Representative images showing localization of Ebf2$^+$/GFP$^+$ cells in relation to expression of NG2 (**D**) and α-smooth muscle actin (α-SMA) (**E**) in skin. Red arrow indicates a Ebf2$^+$ cell without α-SMA expression. Scale bars in are 100 μm (D), 50 μm (E, left) and 10 μm (enlarged images in E).

(**F**) Quantification of Ebf2$^+$/GFP$^+$ cells expressing α-SMA and NG2. Data were from 3 mice. Related to Figure 3.

**Supplementary Figure 4 Skin Ebf2$^+$ cells generated Ebf2$^-$ cells *in vivo*.**

(**A**) Representative FACS plot showing the gating of Ebf2/GFP$^+$Tomato$^-$ and Ebf2/GFP$^+$Tomato$^+$ cells within skin stromal cells (CD45$^-$TER119$^-$CD31$^-$) at 3 months after the last tamoxifen injection. Non-transgenic mice were used as controls for background signals of GFP and tomato in samples from Tg *Ebf2-Egfp* x *Cre$^{ERT2}$x tomato* mice.

(**B**) The proportion of the Ebf2$^+$Tomato$^+$ cells within total Ebf2$^+$ cells at 3 months after tamoxifen injection. Each dot represents data from one mouse. Data are from 3 independent experiments and the horizontal bar represents mean.

(**C**) One representative FACS profile showing the Tomato$^+$ cell subsets within total CD140a$^+$SCA1$^+$ (PαS) cells at 3 months after tamoxifen injection.

(**D-E)** Proportion of Ebf2$^+$Tomato$^+$ (**D**) and Ebf2$^-$Tomato$^+$ cells (**E**) within total PαS MPCs. ns, no significant difference by unpaired *t*-test in (**D**). ** P<0.01, by unpaired Mann-Whitney test (**E**).

(**F-H**) Localization of the Tomato$^+$ cells in relation to NG2 (**F**), NESTIN (**G**) α-SMA expression (**H**) at 3 months after tamoxifen injection. The panorama image was presented as maximum intensity projection and a magnified area was shown as single Z-stack image. Scale bars are 50 μm (F,G and H left) and 10 μm (enlarged images in H).

(**I**) The fractions of Tomato$^+$ cells expressing NG2, NESTIN and α-SMA at 3 months after tamoxifen injection.
Related to Figure 3.

**Supplementary Figure 5 Skin Ebf2$^+$ and Ebf2$^-$PαS MPCs exhibit similar hematopoiesis supportive function to BM MSCs**.
(**A**) The experimental layout. Lin$^-$SCA1$^+$KIT$^+$ (LSK) CD150$^+$ (HSCs) were used for the co-cultures with skin MPCs and BM MSC subsets. The cells were analyzed phenotypically at 3 days after the co-culture.
(**B**) Phenotypical analysis of LSKCD150$^+$ HSCs after 3-day co-culture with either skin Ebf2$^+$ (circle) or Ebf2$^-$PαS (square) MPCs or BM MSCs.
(**C**) CFU-Cs from LSKCD150$^+$ HSCs after being cultured with skin or BM MSCs for 3 days. CFU-granulocyte-macrophage (GM) and CFU-granulocyte, monocyte and erythrocyte (GME) were scored at day 10. The CFU-Cs from fresh sorted LSKCD150$^+$ HSCs were presented as condition controls. ns, no significant difference by unpaired *t*-test.
(**D**) The experimental setup for CAFC assay.
(**E**) Representative images of the CAFCs from the LSK cells co-cultured with the MSCs. Scale bars are 100 μm.
(**F**) The numbers of CAFCs after 7-14 days of co-culture. Data are presented as mean ± SEM from 3-5 independent experiments. Each dot represents mean of replicate measurements in each experiment and the horizontal bars represent median values. * $P<0.05$, determined by unpaired *t*-test was used for statistical analysis.
Related to Figure 4

#### Supplementary Figure 1

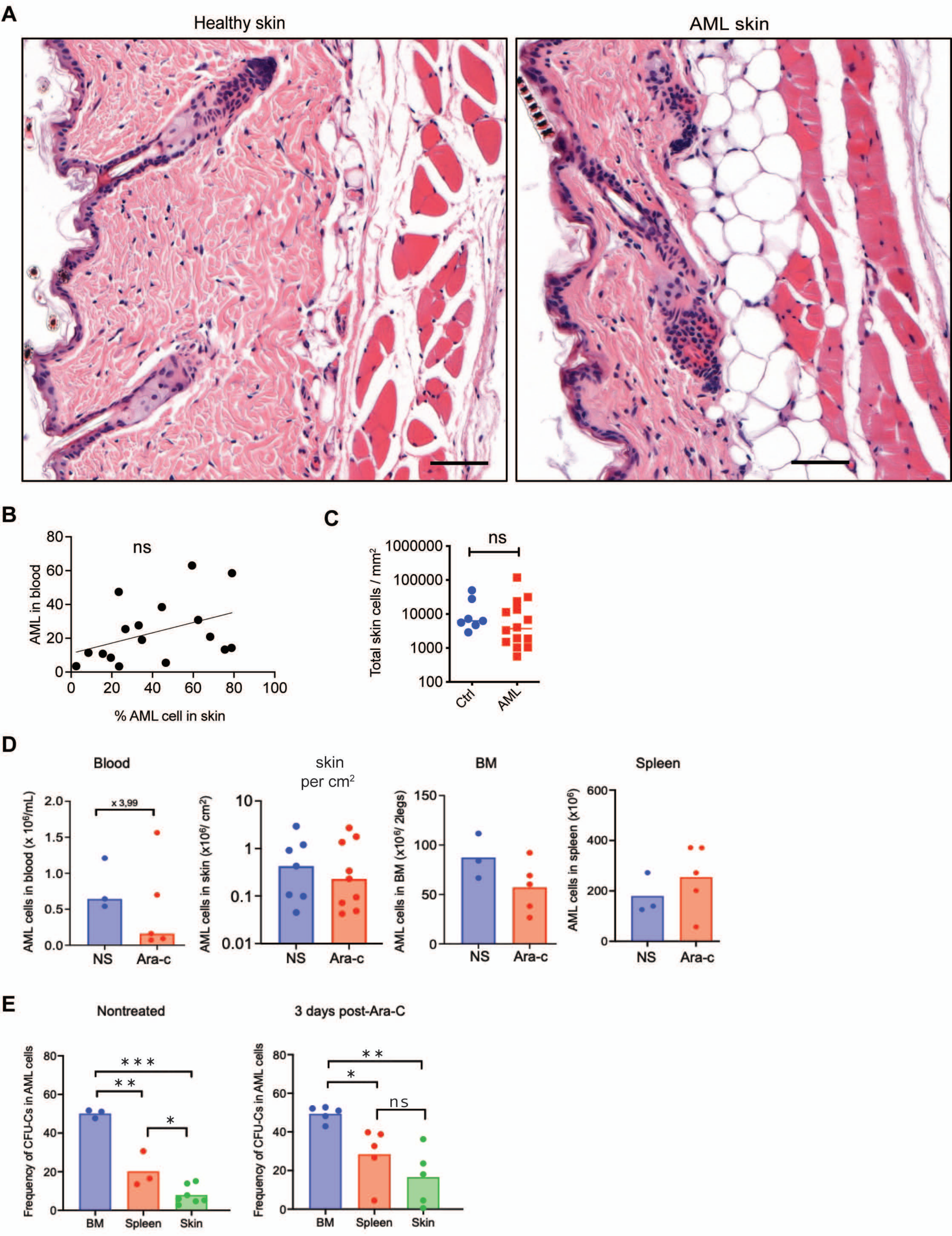

#### Supplementary Figure 2

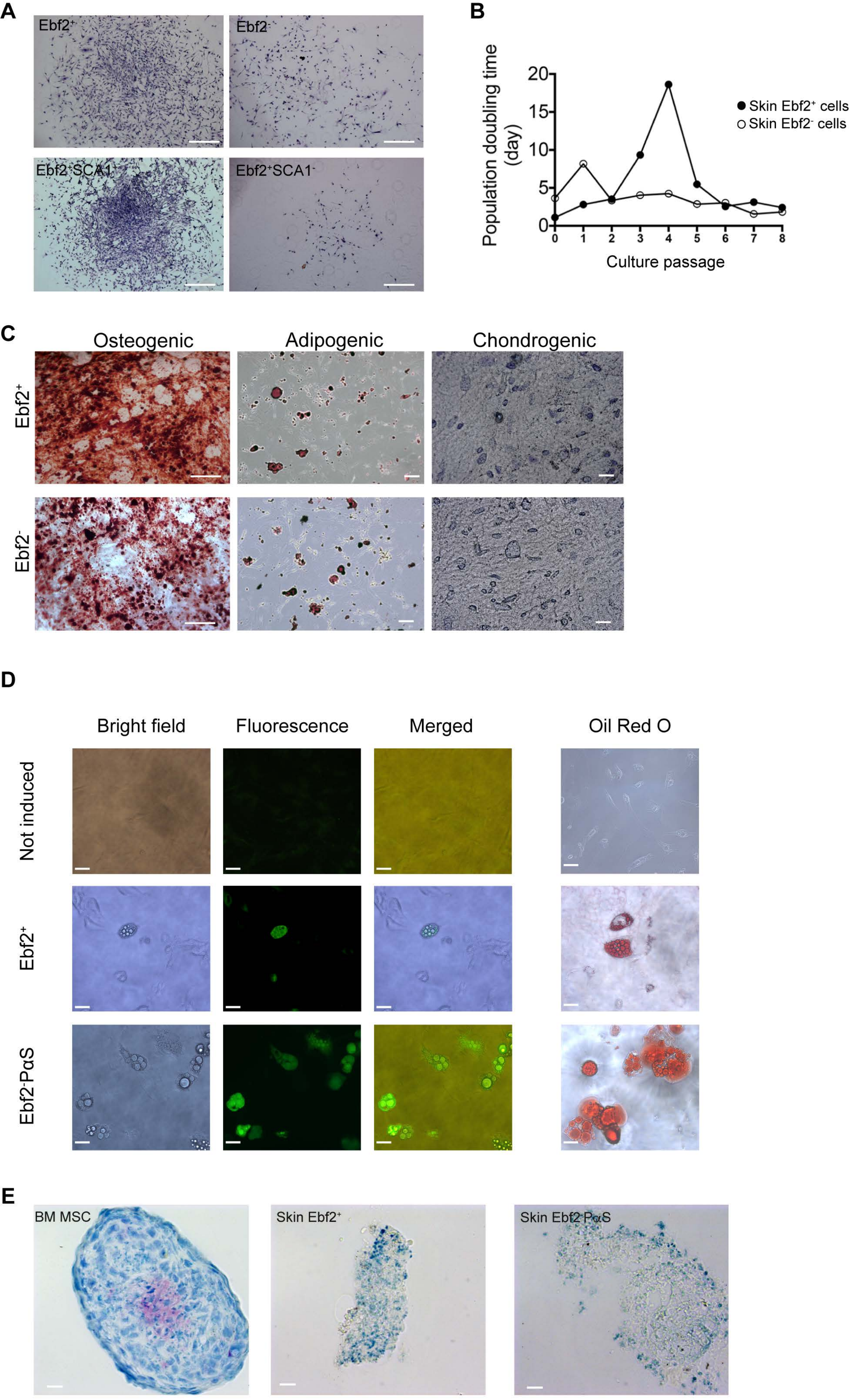

**Supplementary Figure 3**

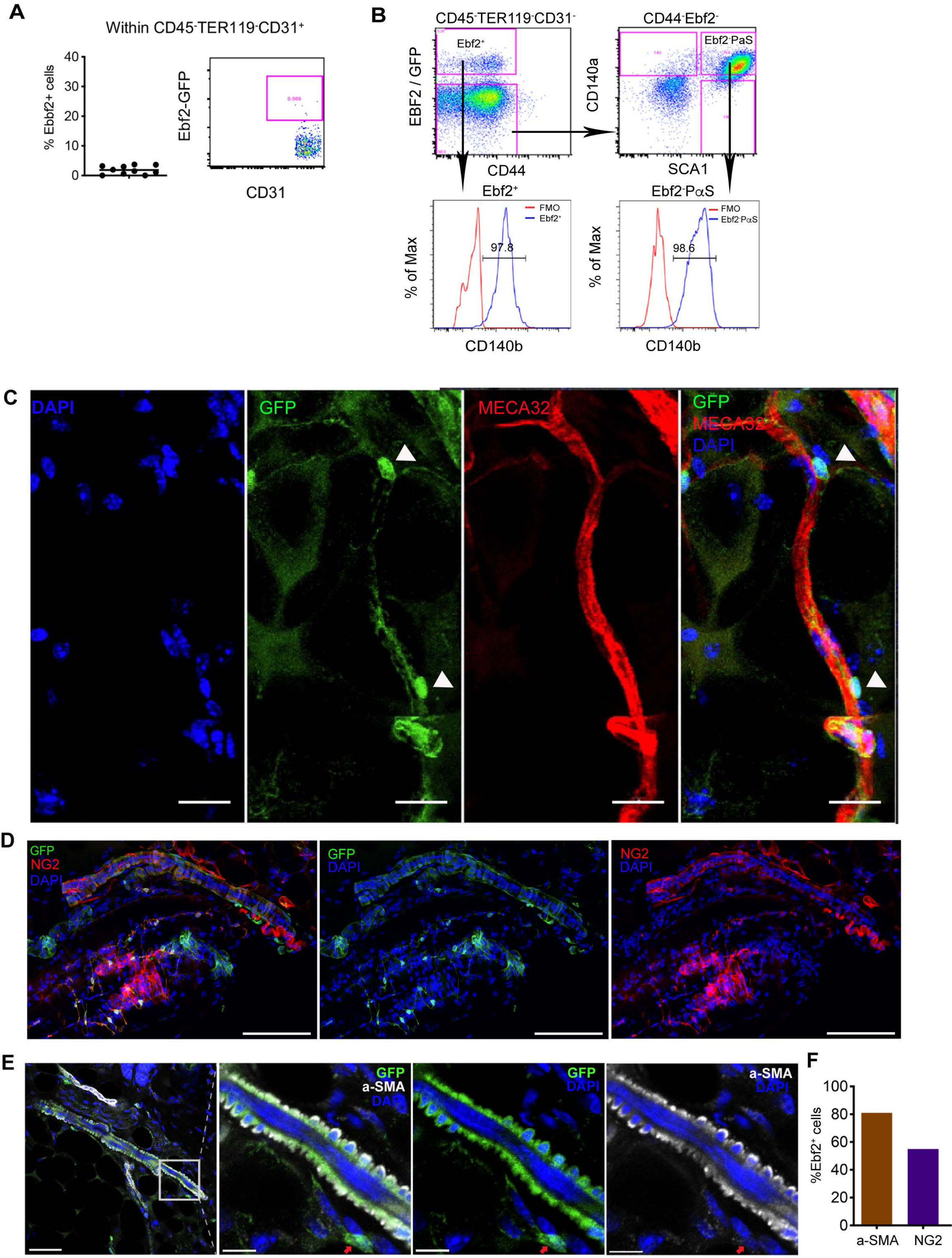

### Supplementary Figure 4

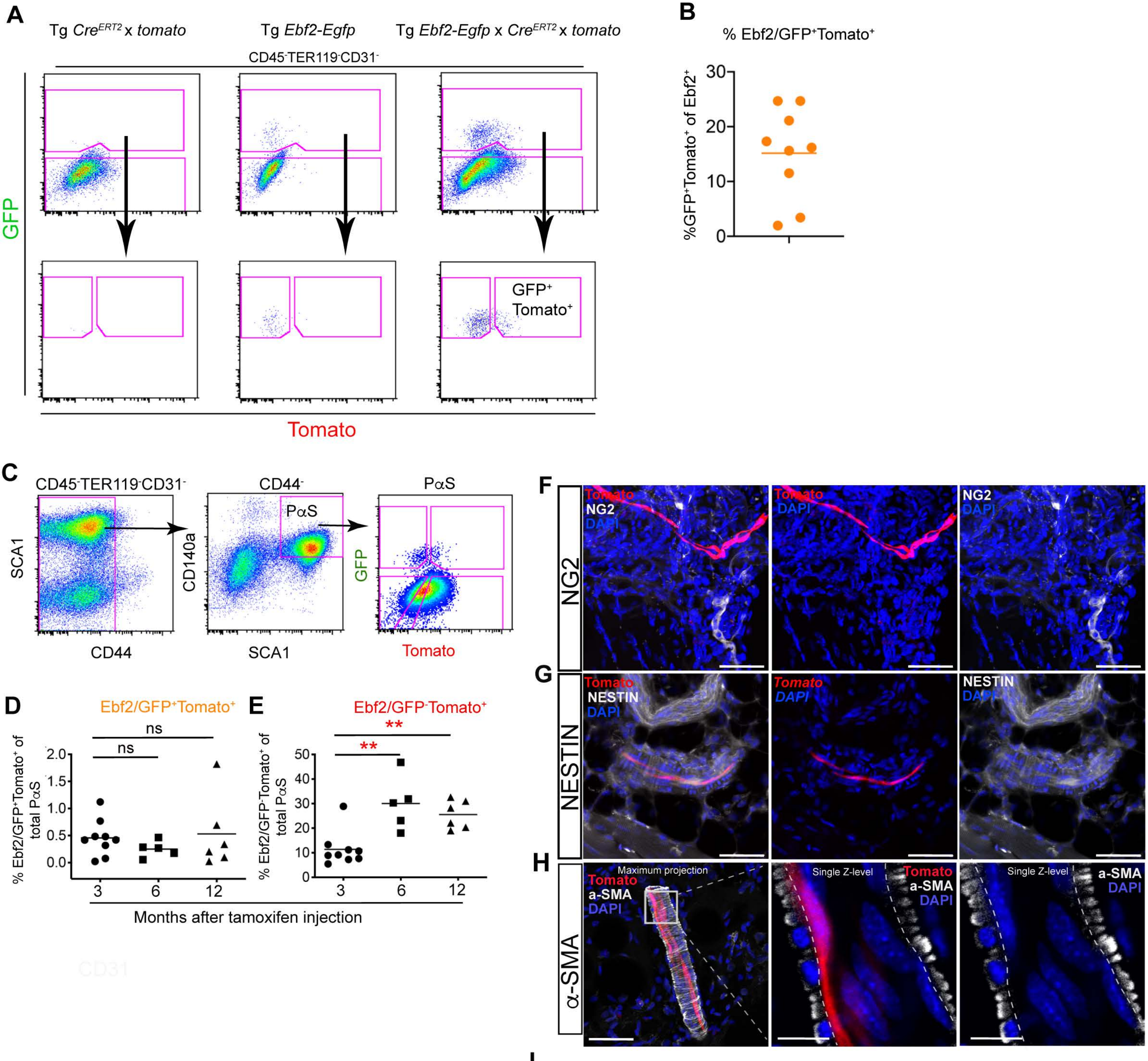


I

| % GFP-Tomato+ cells | | | |
|---|---|---|---|
| CD31 | NG2 | α-SMA | NESTIN |
| 0 | 0 | 28.00 | 18.5 |

### Supplementary Figure 5

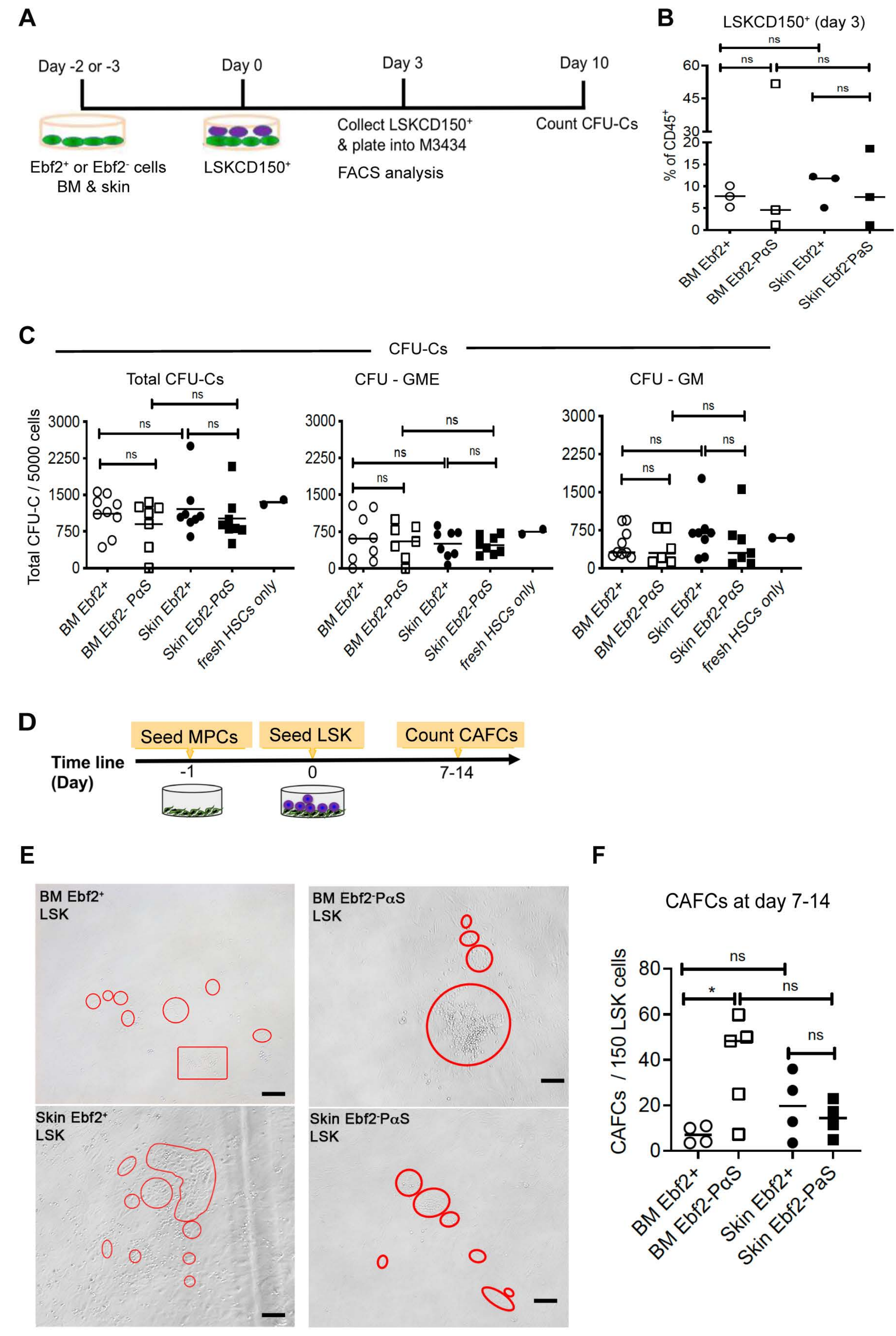
